# Mechanical Instability of Adherens Junctions Overrides Intrinsic Quiescence of Hair Follicle Stem Cells

**DOI:** 10.1101/2020.01.17.909937

**Authors:** Ritusree Biswas, Avinanda Banerjee, Sergio Lembo, Zhihai Zhao, Vairavan Lakshmanan, Manando Nakasaki, Vassily Kutyavin, Graham Wright, Dasaradhi Palakodeti, Robert Ross, Colin Jamora, Valeri Vasioukhin, Yan Jie, Srikala Raghavan

## Abstract

Vinculin, a mechanotransducer associated with both adherens junctions (AJ) and focal adhesions (FA) plays a central role in force transmission through these cell-cell and cell-substratum contacts. Here we describe the conditional knock out (KO) of vinculin in murine skin. Remarkably, we find that the loss of vinculin function results in the loss of bulge stem cell (BuSC) quiescence. We demonstrate that vinculin KO cells are impaired in force generation resulting in mechanically weak AJs. Mechanistically, vinculin functions by keeping α-catenin in a stretched conformation, which in turn regulates the retention of YAP1, another potent mechanotransducer and regulator of cell proliferation, to the junctions. Conditional KO of α-catenin specifically in the BuSCs further corroborates the importance of stable AJs in the maintenance of quiescence and stemness. Altogether, our data provides definitive mechanistic insights into the hitherto unexplored regulatory link between the mechanical stability of cell-junctions and the maintenance of BuSC quiescence.

## Introduction

Stem cells are confined to niches that provide them with the appropriate microenvironment to regulate their proliferative and regenerative potential (Nowak et al., 2008; Pennings et al., 2018). In the mammalian skin, stem cells reside in a specialized niche known as the bulge, located at the lowest permanent portion of the hair follicle and are referred to as the hair follicle stem cells (HFSCs) or bulge stem cells (BuSCs) (Blanpain et al., 2004; Liu et al., 2003; Morris et al., 2004; Tumbar et al., 2004). Hair follicles undergo a cycle of growth (anagen), regression (catagen) and rest (telogen) with the first two cycles being synchronized. A defining feature of the BuSCs is that they are relatively quiescent, but their activation, which occurs in a regulated manner, contributes to hair follicle growth during anagen. After a short proliferative burst, the stem cells move into quiescence until the next hair cycle. Thus, maintaining the BuSCs in the quiescent state is critical for the regulation of hair follicle cycling. The BuSCs are maintained in their niche through cell-cell and cell-substratum interactions, as well as cell intrinsic factors (transcription factors, microRNAs and epigenetic regulators) and secreted factors (Wnts, BMPs, etc) (Gattazzo et al., 2014; Wang and Wagers, 2011). While there is a growing list of cell-intrinsic factors and signaling molecules that regulate the quiescence of the BuSCs, the role of cell-cell adhesion, such as those mediated through AJs, remains underexplored.

AJs in skin are nucleated through homotypic interactions between E-cadherin on neighboring cells (Rübsam et al., 2018) and remodeling of the junctions under tension has been shown to be one of the key regulator of epithelial morphogenesis (Pinheiro and Bellaïche, 2018). Interestingly, AJs of quiescent bulge stem cells express high levels of E-cadherin while reduced levels of E-cadherin in anagen is associated with increased cell proliferation (Lay et al., 2016), suggesting an inverse correlation between junctional stability and proliferation. The core adhesive junctional complex is formed by the interaction of intracellular domain of E-cadherin with an array of catenins including p120, β-catenin and α-catenin (Niessen and Gottardi, 2008). The reinforcement of the junction is achieved by actinomyosin generated force-dependent recruitment of vinculin through its interaction with α-catenin to form mature AJs (Imamura et al., 1999; Le Duc et al., 2010; Miyake et al., 2006; Sumida et al., 2011; Watabe-Uchida et al., 1998; Weiss et al., 1998; Yonemura et al., 2010). Vinculin is a mechanotranducer that is associated both with cell-substratum and cell-cell adhesions. Vinculin comprises a head and tail domain, separated by a linker region. The interaction between the head and tail domain, renders vinculin inactive, and its unfolding, which happens in a force-dependent manner, creates docking sites for several actin-remodeling enzymes (Bays and DeMali, 2017). The Head domain of vinculin binds to α-catenin at the vinculin-binding site (VBS). α-catenin, like vinculin, can also exist in an auto-inhibited state and it is estimated that its unfolding requires ∼5pN of force at a force loading rate of a few pN per second, which then unmasks the binding site for vinculin (Pang et al., 2019; Yao et al., 2014),.

The high-affinity binding of vinculin to α-catenin (Pang et al., 2019; Yao et al., 2014) in turn potentially keeps α-catenin in an open conformation, releasing the vinculin tail domain for binding to actin. The vinculin-actin interface can withstand a force of ∼ 10 pN for ∼10 s (Huang et al., 2017), whereas the vinculin-VBS interface on α-catenin can withstand forces of ∼ 10 pN for more than 1000s (Le et al., 2019). Therefore, vinculin establishes a stable force-transmission pathway between α-catenin and actin and likely interacts with numerous vinculin-binding factors in a force-dependent manner. This stabilizes the protein-protein interfaces of α-catenin—actin and α-catenin—β-catenin. Together α-catenin and vinculin play a central role in transmitting forces to the actomyosin complex, and that is believed to be critical to strengthening the adherens junctions.

Knockout of α-catenin in the skin epidermis results in perinatal lethality due to loss of AJs in the skin, and is associated with the development of precancerous lesions (Vasioukhin et al., 2001). The complete loss of vinculin or absence of vinculin-actin interaction, causes the cells to become less stiff, exert lower traction forces, and unable to reorganize their cytoskeleton (Alenghat et al., 2000; Huveneers et al., 2012; Le Duc et al., 2010; Ohmori et al., 2010). Global ablation of vinculin results in embryonic lethality by E10. The primary defect is impairment in the development of the nervous system and heart (Xu et al., 1998). The conditional KO of vinculin in cardiomyocytes results in animals that develop dilated cardiomyopathy, which is attributed to defective cell-cell and cell-matrix junctions. However, the role of vinculin in the skin, particularly in maintaining the BuSC niche has not been explored. In this study, by employing a skin-specific conditional KO, we demonstrate a hitherto unknown function of vinculin in maintaining stem cell quiescence by regulating junctional stability. Mechanistically we show that the binding of vinculin to α-catenin keeps it in an open/stretched conformation, which in turn sequesters YAP, a potent activator of the cell cycle to the junctions. Taken together our results provide novel insights into the role of junctional proteins in maintaining stem cell quiescence.

## Results

### BuSC compartment is associated with robust cell-cell junctions

We investigated the correlation between junctional stability and quiescence during hair follicle cycling. At the start of a new hair cycle; the telogen/anagen (T/A) transition (WT postnatal day 22, (P22) we observed a reduction in the expression of E-cadherin at the growing front of the hair follicle associated with a weak cortical actin network and Ki67 positive proliferating cells (Fig 1 A). At the telogen stage (P52), we observed strong expression of E-cadherin at the AJs (as previously reported, (Lay et al., 2016)) associated with a thick cortical actin network and a loss of Ki67 positive cells (Fig 1B). Electron micrograph (EM) images of the P24 T/A follicles and P54 telogen follicles corroborated this observation. In the P24 T/A follicles (Fig 1C), we observed a reduction in the number of desmosomes (asterisks in Fig 1C’) at the growing tip of the follicle. On the other hand the P54 bulge stem cells (Fig 1D) were associated with numerous desmosomes (asterisks in Fig 1D’). Since desmosomes are known to be associated with AJs we generated a high magnification EM images of the junction between the bulge stem cells in a P54 follicle. As seen in Fig 1E, the electron-dense desmosomes were closely associated with AJs (schematic, Fig 1E). These data suggested to us that the increased number of desmosomes in the P54 telogen bulge was a proxy for increased number of AJs. We next looked at the expression of vinculin and α-catenin, two important AJ proteins associated with the actin network, in the P22 T/A follicles and in the bulge compartment of a P54 telogen follicle. There was a strong junctional expression of α-catenin and vinculin in the BuSCs at P54 (Fig 1 E-G) in contrast to the growing front of the P22 T/A follicle that was associated with reduced expression of α-catenin and vinculin (Fig S1A, B). Since subtype cadherin switching between E-cadherin and P-cadherin plays an important role in hair follicle cycling (Hirai et al., 1989; Müller-Röver et al., 1999), we looked at the P-cadherin expression during hair cycling. We observed increased P-cadherin expression in the growing front of the P22 T/A follicle, corresponding to the area of low E-cadherin expression (Fig S1C). In contrast, the expression of P-cadherin was absent from the bulge compartment of P52 telogen follicles, and was confined to the secondary hair germ (Fig S1D). EM analysis of the secondary hair germ (Fig S1E) revealed that the number of desmosomes was markedly reduced between the hair germ cells compared to those in the overlying bulge stem cell (asterisks in Fig S1E’). Taken together, these data suggested that the quiescent bulge cells have more cell-cell junctions compared to growing follicles, prompting us to ask what role these junctions may play in maintaining the stem cell compartment.

**Figure 1:**
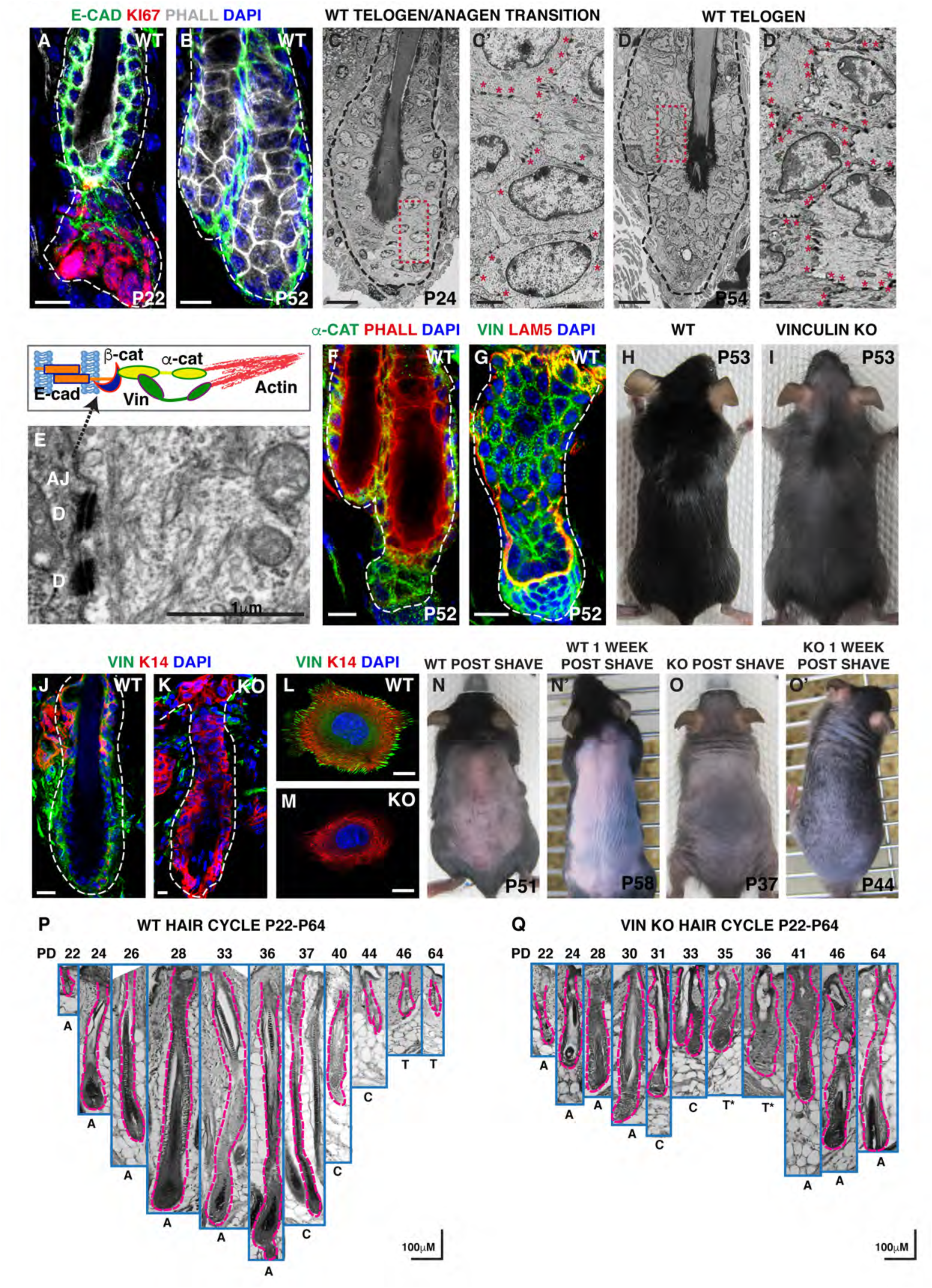
Status of cell-cell junction proteins in hair follicles and phenotype of vinculin KO in skin epidermis: WT Telogen-Anagen (T/A) transition (P-22) and Telogen (P-52) follicles labeled with E-cadherin (green), Ki67 (red), phalloidin (gray) and DAPI (blue) in (A, B). Scanning electron micrograph (SEM) of the WT cell-cell junctions at T/A transition (P-24) and Telogen (P-54) follicles (C, D). Insets: high mag images of the areas indicated by the red dashed box (C’, D’). Red asterisks mark desmosomes. High mag SEM of cell-cell junction indicating desmosomes (D) and adherens junctions (AJs) and schematic of different components of the AJ (E). WT (P-52) junctions labeled with α-catenin (green), phalloidin (red) and DAPI (blue) and vinculin (green) laminin 5 (red) DAPI (blue) (F, G). Phenotype of WT and vinculin conditional KO animals (H, I). WT and vinculin KO hair follicles and keratinocytes labeled with vinculin (green), K14 (red) and DAPI (blue) (J-M). WT and vinculin KO animals analyzed for hair regrowth one-week after shaving (N’, O’). Images taken immediately after shaving (N, O). Histological analysis of hair follicles from WT and vinculin KO animals from P-22-P-64 (P, Q). Scale bar: 10μM (A, B, C, D, F, G and J-M), 2μM (C’-D’), 1μM (E)100μM (P, Q).

### Conditional deletion of vinculin in skin epidermis results in loss of telogen phase

To explore the function of the AJ protein vinculin in maintaining the stem cell niche, specifically its purported role in regulating acto-myosin force-driven junctional reinforcement, we generated the conditional KO of vinculin in the skin epidermis by crossing the vinculin floxed mice with the K14-Cre mice. The vinculin KO mice were born in the correct Mendelian ratio. However they displayed a thinning hair coat, first visible at postnatal day 9 (P9), which got progressively worse as the animals aged (Fig 1H, I). Interestingly upon aging these animals did not lose all their pellage HFs, but displayed a severely thinned hair coat (chased for 1 year, Extended Supplementary Fig1A, B). The loss of vinculin from the skin epidermis and isolated keratinocytes was validated through indirect immunofluorescence and immunoblot analysis (Fig1J-M, Fig S1F). To better understand the HF phenotype, we took WT and KO animals from P1-P64, isolated the anterior back skin, embedded these in paraffin, sectioned and performed Hematoxylin and Eosin (H&E) staining on these sections. We analyzed the morphology and size of the HFs during the first (P1-P21) and second (P22-P64) hair cycles. Histological analysis of WT and KOs revealed that the HFs in the vinculin KO animals were much shorter, deformed and cycling significantly faster than those in the WT (Fig 1P, Q and Fig S1G, H). In WT HFs, morphogenesis ends at P15 followed by catagen (P16-P19) and telogen (P20-22). By comparison, the KO HFs enter catagen at P10, and have a slightly shorter telogen phase (P20-21) (Fig S1G, H). The second hair cycle in WT animals is marked by a growth phase for 12 days (P24-36) followed by catagen for 8 days (P37-44) and a long telogen period lasting approximately eight weeks (Fig 1J). By comparison, in the vinculin KO animals the second anagen lasts about 9 days (P22-P30), followed by a short catagen (P31-P33) and a telogen-like stage (P35-P37). Subsequently the HFs became asynchronized suggesting that in the vinculin KO the follicles undergo the entire second hair cycle by P37 (Fig S1G). The transient telogen phase in the KO follicles was further corroborated by performing shaving experiments during the second telogen (WT P51 and KO P37). The KO follicles grew back hair one-week post-shaving suggesting absence of a prolonged resting phase as compared to their wild type counterparts (Fig 1N-O’). Taken together these data suggest that the absence of vinculin results in the loss of the telogen (resting) phase (Fig S1I).

### Expansion of bulge stem cell compartment and loss of quiescence in vinculin KO follicles

The formation of a club hair follicle occurs at the onset of the long second telogen stage (P46-P110). The club hair is associated with the old bulge, while a new bulge is formed by stem cells that escape apoptosis during the destructive (catagen) phase of the hair cycle (Hsu et al., 2011). The presence of the club hair results in a thicker hair coat in mice. In the vinculin KO animals while the first telogen follicles appeared histologically similar to WT follicles, we noticed the complete absence of club hair follicles at the second telogen stage (Fig 2A. A’, B, B’). Since the HFSC of the new bulge are the only contributors to hair regeneration (Hsu et al., 2011; Lay et al., 2016), we characterized the BuSC compartment in the KO follicles. Due to the accelerated HF cycling in the KO we compared follicles at similar morphological stages of hair cycling. For the first telogen we compared WT P21 animals with KO P20. For the second telogen we compared WT animals between P49 to P53, with KO animals from P35 to P37 that morphologically appeared to be telogen follicles. To assess the status of long-term label retaining stem cells (LRCs) of the HFs we performed ethynyldeoxyuridine (EdU) pulse chase experiments. The retention of the EdU label in the bulge is a read out of the presence of the quiescent stem cells (Cotsarelis et al., 1990; Morgner et al., 2015). The expression of EdU was significantly reduced in the first telogen follicles, and completely absent from the second telogen bulge compartment, suggesting a loss of these quiescent pools of stem cells in the vinculin KO animals (Fig 2C, C’, D, D’). On the other hand, and in concordance with the loss of LRCs, there was an increase in proliferation as assessed by the expression of Ki67 in the KO follicles (Fig 2E, E’, F, F’). Given the loss of the long-term LRCs in the KO, we examined the status of the BuSCs by labeling skin sections with other stem and progenitor cell markers CD34, SOX9, K15, LGR5 and LRIG1 (Liu et al., 2003; Schepeler et al., 2014) (Fig S2D). CD34 (cluster differentiation 34) a cell surface protein is a common hematopoietic stem cell marker that also happens to mark the multipotent BuSCs of the HFs. The expression of CD34 was significantly reduced in the BuSCs of the first telogen KO follicles (Fig S2A, A’, S2C) and completely absent in the second telogen follicles (Fig S2B, B’) (Morris et al., 2003). Another highly expressed stem cell marker in the bulge region is K15 (keratin 15) that co-localizes with CD34+ cells. These K15 positive cells from the bulge region have been shown to be multipotent (Morris et al., 2004). The expression domain of K15 in the first and second telogen KO follicles was expanded, such that the entire follicle appeared to be K15 positive. The expression of K15 in WT follicles was largely limited to the bulge region (Fig S2E, E’, F, F’). The transcription factor SOX9 (sex determining region Y box9) is one of the earliest markers of putative BuSCs. The expression of SOX9 is seen as early as E15.5 in developing follicles and the conditional KO of SOX9 from the epidermis results in loss of LRCs and cycling matrix cells, underscoring its role in maintaining the stem cell niche (Nowak et al., 2008; Vidal et al., 2005)). The expression domain of SOX9 was expanded throughout the entire first and second KO telogen follicles, compared to a more restricted expression in the WT (Fig S2I, I’, J, J’). LGR5 (leucine-rich repeat-containing G-protein coupled receptor 5) is an intestinal stem cell marker that is also expressed in the lower bulge region and hair germ of growing follicles (Jaks et al., 2008). In the first telogen follicles the expression of LGR5 was found predominantly in the bulge region and the outer root sheath of the WT follicle (Fig S2G). In comparison, the expression of LGR5 was expanded in the KO follicles (Fig S2G’). In the second telogen follicles, the expression of LGR5 was restricted to the secondary hair germ in WT follicles (Fig S2H) but was expressed throughout the follicle in the KOs (Fig S2H’). LRIG1 (Leucine-rich repeats and immunoglobulin-like domain protein 1) is a stem cell marker that is expressed in the junctional zone which contains actively cycling cells (Jensen et al., 2009; Page et al., 2013). LRIG1+ cells are devoid of canonical BuSC markers like CD34 and LGR5, however these cells can give rise to all epidermal lineages in reconstitution assays (Jensen et al., 2009; Page et al., 2013). The expression of LRIG1 was restricted to the junctional zones of the WT first and second telogen follicles (Fig S2K, L) and the expression domain was expanded beyond the junctional zone in the KO follicles (Fig S2K’, L’). Taken together our analysis of the stem cell compartment suggests that stem cell homeostasis is severely compromised in vinculin KO animals. In addition to the loss of LRCs, other stem cell or progenitor cell markers are either not expressed or misexpressed, suggesting premature entry of stem-like cells into the cell cycle. To explore whether these cells in the bulge region do indeed behave like the canonical BuSCs, we performed hair and skin reconstitution experiments with WT and vinculin KO P0 keratinocytes and fibroblasts. While the WT cells were able to contribute to the formation of hair and skin as judged by the presence of pigmented hair on the back of albino SCID mice, and by histological analyses (Fig 2G, H), the vinculin KO cells failed to contribute towards the formation of hair and skin. Instead we observed the presence of large cyst-like structures, suggesting compromised differentiation of hair follicle cells (Fig 2I, J).

**Figure 2:**
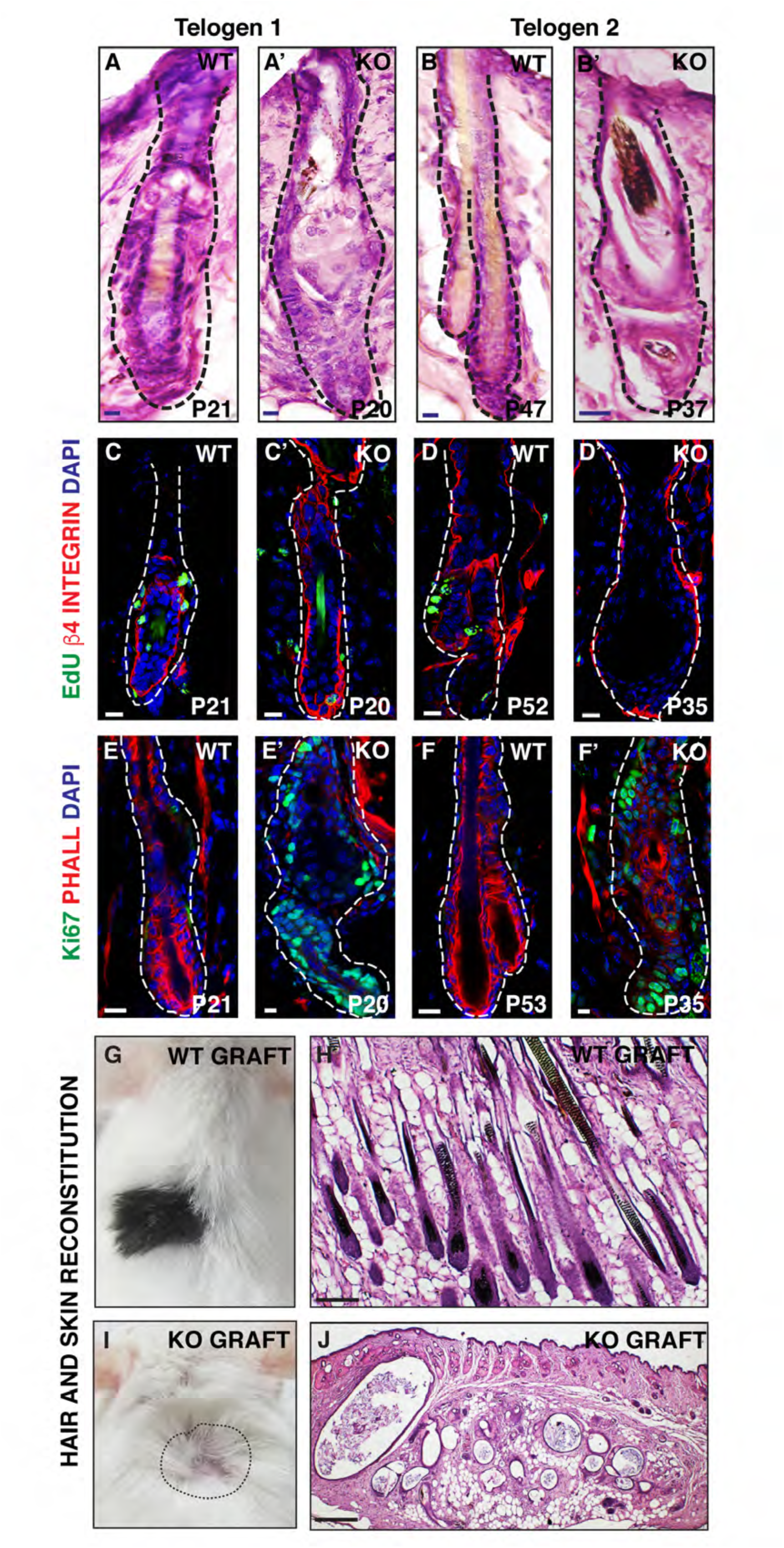
Analysis of stem cells in vinculin KO mice: Histological analysis of hair follicles from WT (P-21, P-46) and vinculin KO (P-20, P-37) animals (A, A’, B, B’). WT and vinculin KO follicles LRCs labeled with the CLIK-iTEdU Fluor488 (green), β4 integrin (red) and DAPI (blue) (C, C’, D, D’). WT and vinculin KO skin sections labeled with Ki67 (green), phalloidin (red) and DAPI (blue) (E, E’, F, F’). Hair and skin reconstitution performed with WT and KO keratinocytes grafted in NOD SCID mice (G, I), Histological analysis of the WT and KO skin grafts (H, J). Scale bar: 10μM (A-F’), 100μM (H, J).

### Perturbed hair follicle differentiation in vinculin KOs

Given that the stem cell homeostasis was severely perturbed in the vinculin KO animals, we explored whether the establishment of the different layers of the HFs were affected in KO animals, both during early morphogenesis (P6) and in anagen (P30) (Extended Supplementary Fig 2B, B’) (Taylor et al., 2000) Hair morphogenesis initiates through epithelial-mesenchymal interactions during embryonic development and is followed post-natally by the establishment of a pool of transiently amplifying matrix cells that maintain close contact with the underlying mesenchymally-derived dermal papilla (DP)(Oliver and Jahoda, 1988). Upward movement of LEF1 (Lymphoid enhancer-binding factor 1) expressing matrix cells establishes the cortex and the inner root sheath (IRS), which are marked by AE13 (Alpha-esterase-13) and AE15 (trichohyalin) respectively (DasGupta and Fuchs, 1999; Kobielak et al., 2003; Manabe et al., 1996) (Extended Supplementary Fig 2A). The IRS is surrounded by a companion cell layer and the outer root sheath (ORS) that are marked by K6 (keratin 6) and K5 (keratin 5) respectively. We found no discernible differences in the expression patterns of K5 and K6 between WT and vinculin KO P6 follicles (Extended Supplementary Fig 2C). Similarly the expression pattern of AE15 and AE13 was similar in WT and KO P6 follicles (Extended Supplementary Fig 2D, E). Finally the expression of LEF1 that marks the matrix was also largely unchanged between the WT and P6 KO follicles (Extended Supplementary Fig 2F). We next looked at these markers in P30 follicles. We chose this stage, since both WT and KO follicles are in anagen (Extended Supplementary Fig 2B’). We observed occasional cyst like structures in the KO follicles that was accompanied by perturbed differentiation in the KO (Extended Supplementary Fig 2C’). This was particularly evident when we looked at the expression of AE15 and AE13 in the P30 KO follicles, suggesting that both the IRS and cortex were not correctly specified (Extended Supplementary Fig 2D’, E’). Finally, there was reduced expression of LEF1 in P30 KO follicles (Extended Supplementary Fig 2F’). Taken together, these data suggest that while early hair morphogenesis was not perturbed in the vinculin KO animals, loss of vinculin results in defective hair follicle differentiation during later adult hair cycling.

### Loss of vinculin leads to impaired adherens junction formation

Since vinculin is expressed both at the cell-cell AJs and the cell-substratum FAs, we investigated the status of these junctions in the BuSC compartment in the first and second telogen follicles. We labeled WT and vinculin KO HFs with E-cadherin and α-catenin antibodies. In the WT P21 and P56 follicles the expression of E-cadherin was limited to the AJs (Fig 3A, A’ C, C’). In the KO P21 and P36 follicles the expression of E-cadherin was disrupted (Fig 3B, B’, D, D’). Likewise α-catenin was expressed in the AJs of the P21 and P52 WT follicles (Fig 3E, E’, G, G’), and this expression was quite disrupted in the KO P20 and P36 follicles with expression not being limited to the cell junctions (Fig 3F, F’ H, H’). We next examined the ultrastructure of the AJs, by processing both WT and KO skin sections for scanning electron microscopy (SEM). Like with the immunofluorescence images, we chose to focus on the bulge compartment of first and second telogen follicles. We used the easily identifiable electron dense desmosomes in the electron micrographs as the readout of AJ formation. The organization of normal AJs is a prerequisite for desmosomal junction formation (Lewis et al., 1997). In the WT SEMs we observed evenly spaced desmosomal structures, at cell-cell junctions associated with plasma membranes that were closely apposed (Fig 3I, I’, K, K’). In comparison the number of desmosomes in the KO follicles was significantly reduced (extended data ES2) and this was associated with separation in the plasma membrane between 2 adjacent cells (Fig 3J, J’, L, L’), suggesting that cell-cell junctions in the KO follicles were weak. Interestingly we did not observe a significant decrease in desmosome numbers in the inter follicular epidermis (IFE) (extended supplementary Fig 3) nor the separation between plasma membrane in the IFE suggesting that this junctional instability was primarily limited to the BuSC compartment.

**Figure 3:**
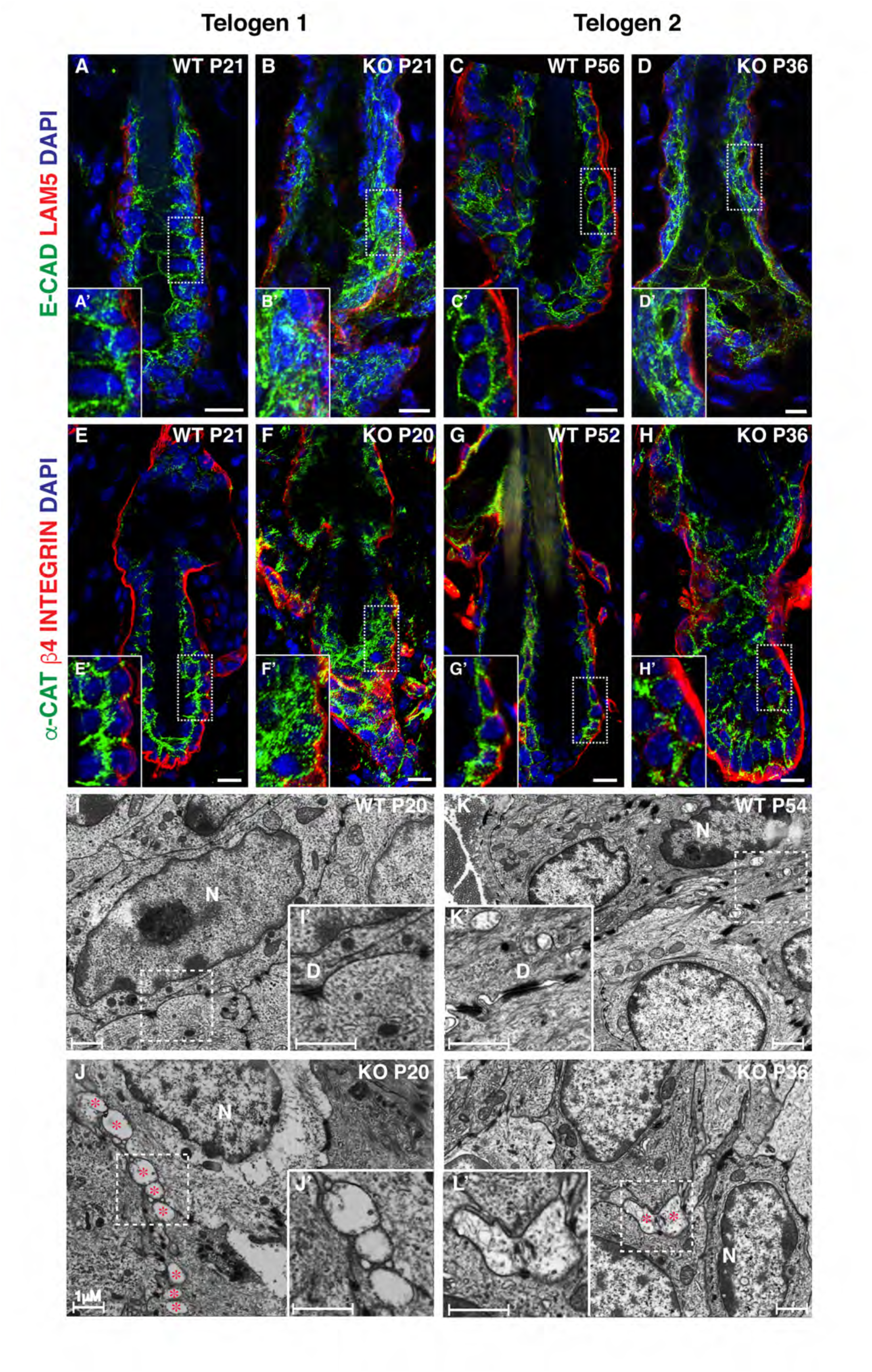
Analysis of adherens junctions of the bulge stem cells at first and second telogen follicles: WT and vinculin KO bulge stem cells junctions labeled with E-cad (green), lam5 (red) and DAPI (green) (A-D), Insets: high mag images of the junctions indicated by the dashed box (A’-D’). WT and vinculin KO bulge stem cells junctions labeled with α-catenin (green), β4-integrin (red) and DAPI (blue) (E-H), Insets: high mag images of the junctions indicated by the dashed box (E’-H’). Electron micrographs of the cell-cell junctions in WT and KO telogen follicles (I-L), Insets: high mag images of the areas indicated by the white dashed box (I’-L’). N-Nucleus, D-Desmosomes, Red asterisks indicate the areas of separation between the cells. Scale bar: 10μM (A-H), 5μM (A’-H’), 1μM (I-L and I’-L’).

To better understand the dynamics of AJ formation, we transfected primary WT and vinculin KO keratinocytes with an E-cadherin GFP construct and imaged the junctions using a spinning disc confocal microscope. Whereas the WT cells were able to form stable junctions in about 3h and sustain these junctions over a long period of time, the KO cells were more dynamic and unable to form stable junctions (movies M1, M2, M3, M4). This phenotype was further validated when we induced the formation of AJs in primary cells using calcium switch assays. After 24h the WT keratinocytes showed a single line of E-cadherin expression at the closed adhesion zippers associated with F actin fibers labeled with phalloidin (Fig S3E). In contrast the KO keratinocytes were unable to zipper up the junctions even 24h after calcium switch (Fig S3F). Phalloidin staining in the KO keratinocytes showed thick bundles of F actin fibers (Fig S3F). Likewise we looked at the expression of α-catenin at the AJs. In WT cells there was punctate expression of α-catenin along the cell junctions (Fig S3G). In contrast, the KO cells displayed elongated structures at the cell-cell junctions, associated with actin stress fibers and a failure to zipper the junction (Fig S3H). The failure to form stable adhesion zippers was further validated by performing SEM on WT and KO cells 24h after calcium switch. As expected in the WT cells, the plasma membranes of the two neighboring cells were closely apposed and associated with thick finger-like filopodial extensions that have been previously described to be essential for the formation of adhesion zippers ((Vasioukhin et al., 2000)) (Fig S3I, I’). In contrast the KO cells displayed thin filipodial extensions, and the plasma membranes of the adjacent cells were not closely apposed (Fig S3J, J’).

Given the disruptions in the AJs, both *in vivo* and in cells in culture, we wanted to ask if the levels of the AJ proteins were changed in the KO compared to the WT. We performed western blot analysis of AJ proteins from both cell and whole skin lysates. Surprisingly the levels of E-cadherin, α-catenin were elevated in both the KO skin and cell lysates compared to the WT with E-cadherin being the most over-expressed. Interestingly the level of β-catenin however was not significantly changed between the WT and KO.

Finally we wanted to address the role of vinculin in cell-substratum junctions and it’s effect on the organization of the basement membrane (BM). We labeled the hair follicles in telogen 2 with anti-collagen IV antibody, and observed that there seemed to be no discernable disruption in the expression of this protein between the WT and vinculin KO follicles (Fig S3A, B). We also used SEM to look at the ultra structural organization of the BM of the IFE. The BM appeared intact, with no appreciable decrease in the number of hemidesmosomes in the KO compared to the WT sample (Fig S3C, D). Taken together, these data suggest that the loss of vinculin specifically affects the stability of the AJs particularly in the hair follicle stem cells.

### Loss of α-catenin in the BuSCs phenocopies the vinculin KO phenotype

Since ablation of vinculin led to the loss of AJ stability, we wanted to explore the role of other AJ proteins in the maintenance of stem cell quiescence. We analyzed the hair follicle cycle and stem cell compartment in the GFAP-Cre-driven α*-*catenin conditional KO mice, where α*-*catenin was depleted in the BuSC compartment, but maintained in the interfollicular epidermis (Silvis et al., 2011). The α*-*catenin KO animals displayed patchy hair growth on their back or complete hair loss. Similar to vinculin KO animals, histological analysis of the back skin of α*-*catenin KO mice revealed defective hair follicle cycling. We observed distorted morphology and size of the hair follicles during the first two hair cycles (P2-P49) (Fig. 4A, B). The hair follicles in P27 α*-*catenin KO mice were morphologically similar to P21 telogen follicles in WT animals. Following the first telogen, the hair follicles never reached the size of the WT anagen follicles (compare P35 WT with P35 KO follicle (Fig. 4A, B)). The delayed entry of the α*-*catenin KO follicles into the telogen phase indicated more actively cycling stem cells. To assess the stem cell compartment, we performed Bromodeoxyuridine (BrdU) pulse chase experiments. We observed significant loss of BrdU at P16 in LRCs in the BuSC compartment, suggesting a loss of quiescence in stem cells in α*-*catenin KO follicles (Fig. S4A-C). The loss of LRCs was associated with increased proliferation as seen by the expression of the proliferation marker Ki67 in the α*-*catenin KO follicles (Fig. 4C, D). We next examined the status of the BuSCs by labeling skin sections at the telogen phase (P21 for WT and P27 for KO) with stem cell markers CD34, SOX9 and K15. CD34 expression was significantly reduced in the bulge compartment in α*-*catenin KO follicles (Fig 4E, F). The loss of expression of CD34 was further validated through FACS analysis of WT and KO BuSCs (Fig. S4D, E). The expression of SOX9 and K15 was expanded throughout the follicles indicating impaired stem cell homeostasis in α*-*catenin KO mice (Fig 4G, H, I, J). LEF1, a transcription factor downstream of Wnt signalling, is crucial for proliferation of the matrix cells in HFs (Rishikaysh et al., 2014). We observed enhanced nuclear localization of LEF1 in α*-*catenin KO HFs, further corroborating increased proliferation in the matrix compartment (Fig. 4K, L).

**Figure 4:**
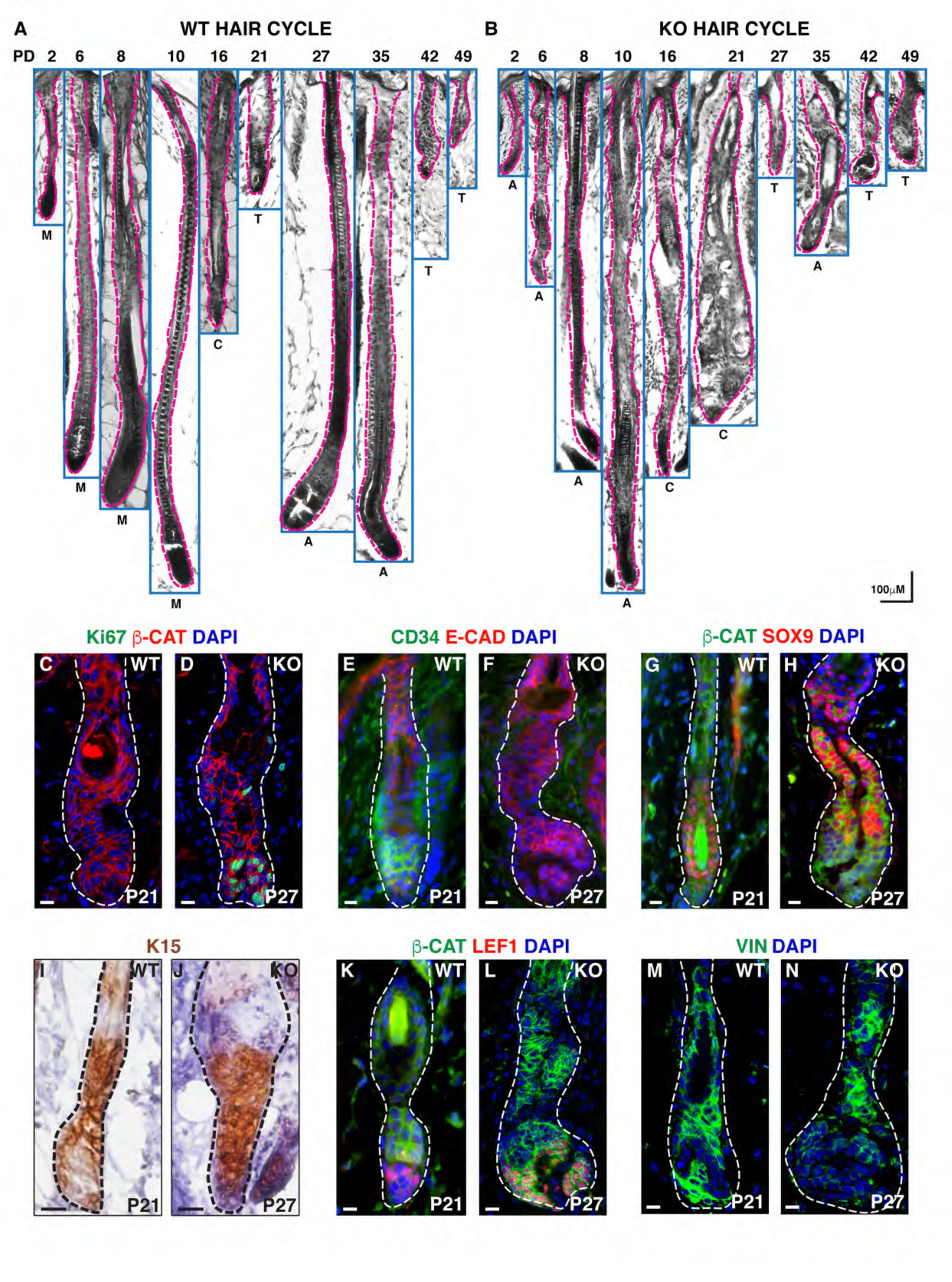
Analysis of the hair cycle and bulge compartment of α-catenin WT and KO telogen follicles. Histological analysis of hair follicles from WT and α-catenin KO animals from first and second hair cycles (A, B). WT and α-catenin KO skin sections labeled with Ki67 (green), β-catenin (red) and DAPI (blue) (C, D). WT and α-catenin KO bulge stem cells labeled with CD34 (green), E-cadherin (red), and DAPI (blue) (E, F). WT and α-catenin KO bulge stem cells labeled with β-catenin (green), Sox-9 (red) and DAPI (blue) (G, H). Immunohistochemistry of WT and vinculin KO bulge stem cells labeled with K15 (I, J). WT and α-catenin KO bulge stem cells labeled with β-catenin (green), Lef1 (red) and DAPI (blue) (K, L). Expression of vinculin (green) in WT and α-catenin KO telogen follicles (M, N). Scale bar: 100μM (A, B), 10μM (C-N).

We analyzed cell junction formation in α*-*catenin KO hair follicles and cultured keratinocytes. As predicted, the localization of E-cadherin in the KO follicles was highly disrupted with most of the protein being located to the cytoplasm and away from cell-cell junctions (Fig. S4F, F’, G, G’). Likewise, in α*-*catenin KO cells in culture, E-cadherin was weakly associated with the AJs (Fig. S4H-K). We next examined the expression of vinculin in the bulge compartment of the α-catenin KO follicles. Compared to the WT P21 follicle, there was a significant reduction in the recruitment of vinculin at the junctions in the α-catenin KO (Fig 4M, N). Finally, loss of α*-*catenin in keratinocytes (Fig S4H, I) also resulted in the failure to recruit vinculin to the AJs and it was localized primarily to the FAs (Fig. S4K).

Taken together these data suggest that the α-catenin KO phenocopies the vinculin KO both in terms of the loss of junctional stability and stem cell quiescence.

### Vinculin provides mechanical stability to AJs

Since loss of both vinculin and α-catenin leads to the formation of unstable AJs, we quantified the forces generated by these junctions. To quantify the strength of the E-cadherin mediated adhesion at AJs (Fig 5A’) in WT, vinculin KO and α-catenin KO cells, we designed a novel microbead displacement experiment as shown in Fig 5A. The force required to destabilize the cadherin-cadherin junctions in vinculin and α-catenin WT keratinocytes was ∼5 to 6nN (Fig 5B, C, D, and G). In comparison, only ∼2.5nN force was sufficient to break the cadherin mediated junctions in the vinculin and α-catenin null keratinocytes (Fig 5B, C, E, H). In both vinculin and α-catenin WT cells there was a strong colocalization of vinculin with α-catenin at the AJ’s (Fig 5D’, G’), whereas in α-catenin KO cells there was an absence of vinculin at the cell-cell junctions, while its localization at the FAs was not affected (Fig 5H’). In the vinculin KO cells on the other hand, there was increased expression of α-catenin at the AJs (Fig 5E’ and Fig S3K) but this seemed insufficient to generate forces similar to WT cells suggesting that the loss of vinculin and α−catenin from the AJs results in a mechanically weak junction that can be easily disrupted when compared to the WT junctions. When we transduced a full-length vinculin construct (VIN-GFP) into vinculin KO keratinocytes we were able to rescue the junctional strength to WT levels (∼6nN) as well as restore the normal expression of α-catenin and vinculin to the AJs (Fig 5B, F, F’). Likewise, when we transfected a full-length α-catenin construct Emerald-α-catenin (EM-α-cat) into α-catenin null keratinocytes the junction strength was significantly rescued as well as the expression of α-catenin and vinculin at the AJs (Fig 5C, J, J’). We would like to note here that the expression levels of the EM-α-catenin construct as well as an α-catenin GFP construct that we used (data not shown) was not as robust as some of the other constructs that we transduced/transfected. That said, the difference between the forces generated by α-catenin WT cells and EM-α-catenin rescued cells was not significant (Fig 5C).

**Figure 5:**
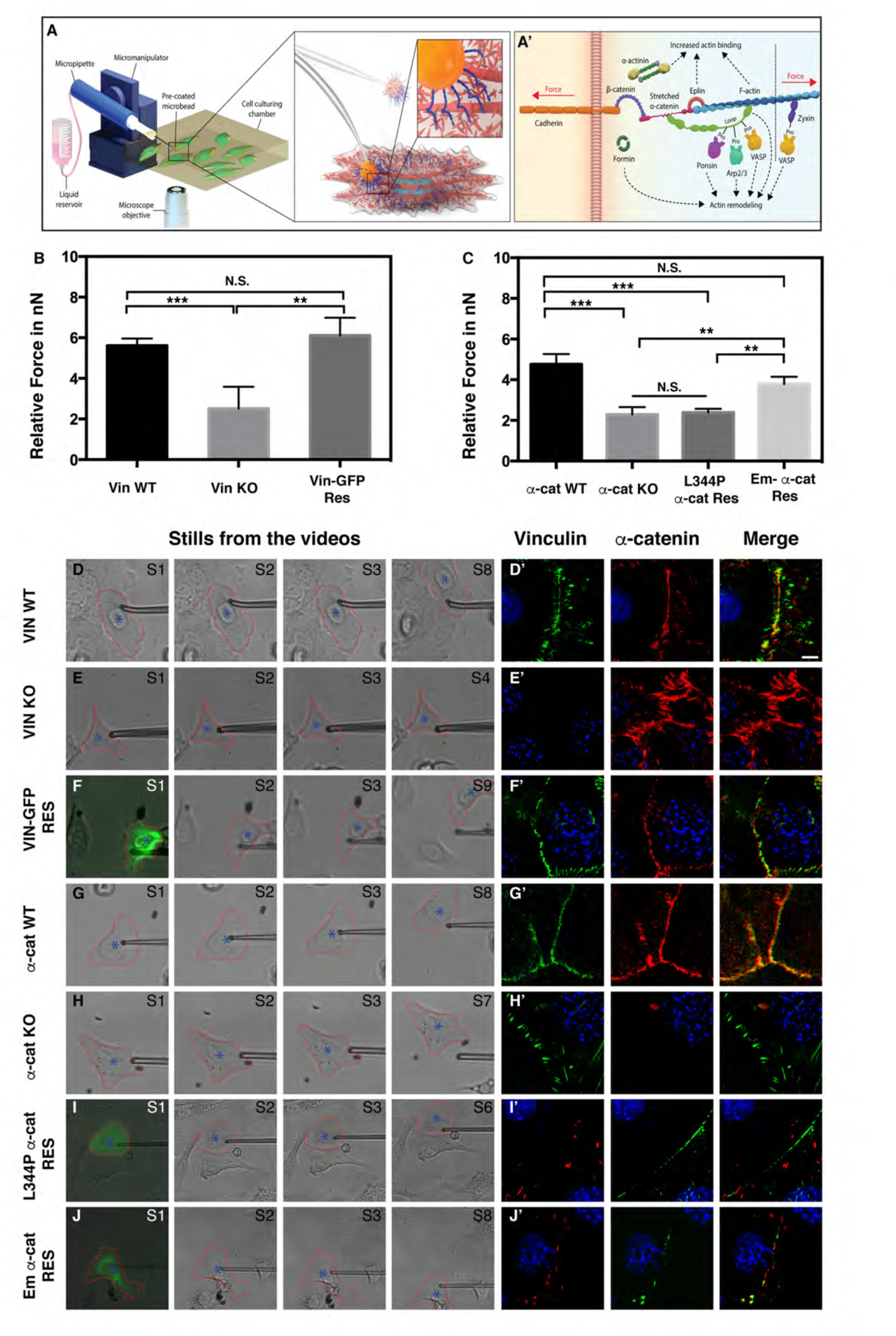
Analysis of force generation by AJs. Schematic representation of the experimental setup, showing the E-cadherin coated bead captured by the glass micro-pipette (A). Schematic of AJ proteins (A’). Histograms representing the relative force measurements required for bead displacement from WT, vinculin-KO, Vinculin-GFP rescued cells (B). Histograms representing the relative force measurements required for bead displacement from WT, α-catenin KO, L344P α-catenin rescued and EM-α-catenin rescued cells (C). Stills of the videos from the bead-detachment experiments performed on vinculin WT (D), vinculin KO (E), vinculin-GFP rescue (F), α-catenin WT (G), α-catenin KO (H), L344P α-catenin rescue (I) and EM-α-catenin rescue cells (J). High magnification images of AJs labeled with vinculin (green), α-catenin (red) and DAPI (blue) from WT (D’), vinculin KO (E’), vinculin-GFP rescue (F’), α-catenin WT (G’), α-catenin KO (H’). High magnification images of cells rescued with L344P α-catenin GFP rescue construct (I) and EM-α-catenin rescue construct labeled for α-catenin (green), vinculin (red) and DAPI (blue). Scale bar: 10μM (D’-J’). \**P* < 0.05, \*\**P* < 0.01, \*\*\**P* < 0.001, N.S. non significant

To address the importance of interaction between vinculin and α-catenin at the AJs, we measured the strength of the AJs in α-catenin KO cells transfected with α-cat-L344P-GFP (α-catenin point mutant that is unable to bind vinculin) (Ming, 2019; Seddiki et al., 2018; Yao et al., 2014). Interestingly, in α-cat-L344P cells the strength of the junctions was around ∼2.5nN, which was similar to the strength of the junction in α-catenin KO cells (Fig 5C, I). Likewise, while the α-cat-L344P-GFP construct localized to the AJs, it failed to recruit vinculin, which was expressed only in FAs (Fig 5I’). This suggested that vinculin played a critical role in reinforcing junctional stability through its interaction with α-catenin and the increased expression of α-catenin in the vinculin KO AJs (Fig 5E’) was unable to compensate for the loss of vinculin. Taken together, these data reveal that vinculin plays a critical role in force generation by the AJs.

### Loss of vinculin affects the conformation of **α**-catenin at AJs and disrupts its interaction with YAP

To correlate the loss of quiescence with junctional instability, we sorted the BuSCs from first telogen WT P-21 and KO P-20 animals using the cell surface markers α6 and CD34 and analyzed its transcriptome. There were markedly fewer α6^hi^/ CD34^hi^ BuSCs isolated from vinculin KO animals compared to WT (Fig S2C). The differentially expressed genes in vinculin KO BuSCs were categorized based on their involvement in specific biological processes (supplementary tables 1-3). We observed upregulation of transcripts of genes involved in the cell cycle, negative regulation of cell death, epithelial cell differentiation, biological adhesion, cell-cell signaling, and cytoskeleton organization (Fig 6A and supplementary table 3). Interestingly, targets of YAP such as *Cdc20, Cdc25b, Ccnb1, Birc5, Tnc, Itga3, Krt7, Krt79, Fzd1* and *Lmna* were upregulated in the KO BuSCs. We validated specific YAP target genes that were involved in cell cycle regulation through real time qPCR analysis (Fig 6B). Pathways that were down regulated involved biological processes such as lipid metabolism, chromatin organization, intracellular transport, regulation of cell development and apoptotic processes (Fig S6J and supplementary table 3).

**Figure 6:**
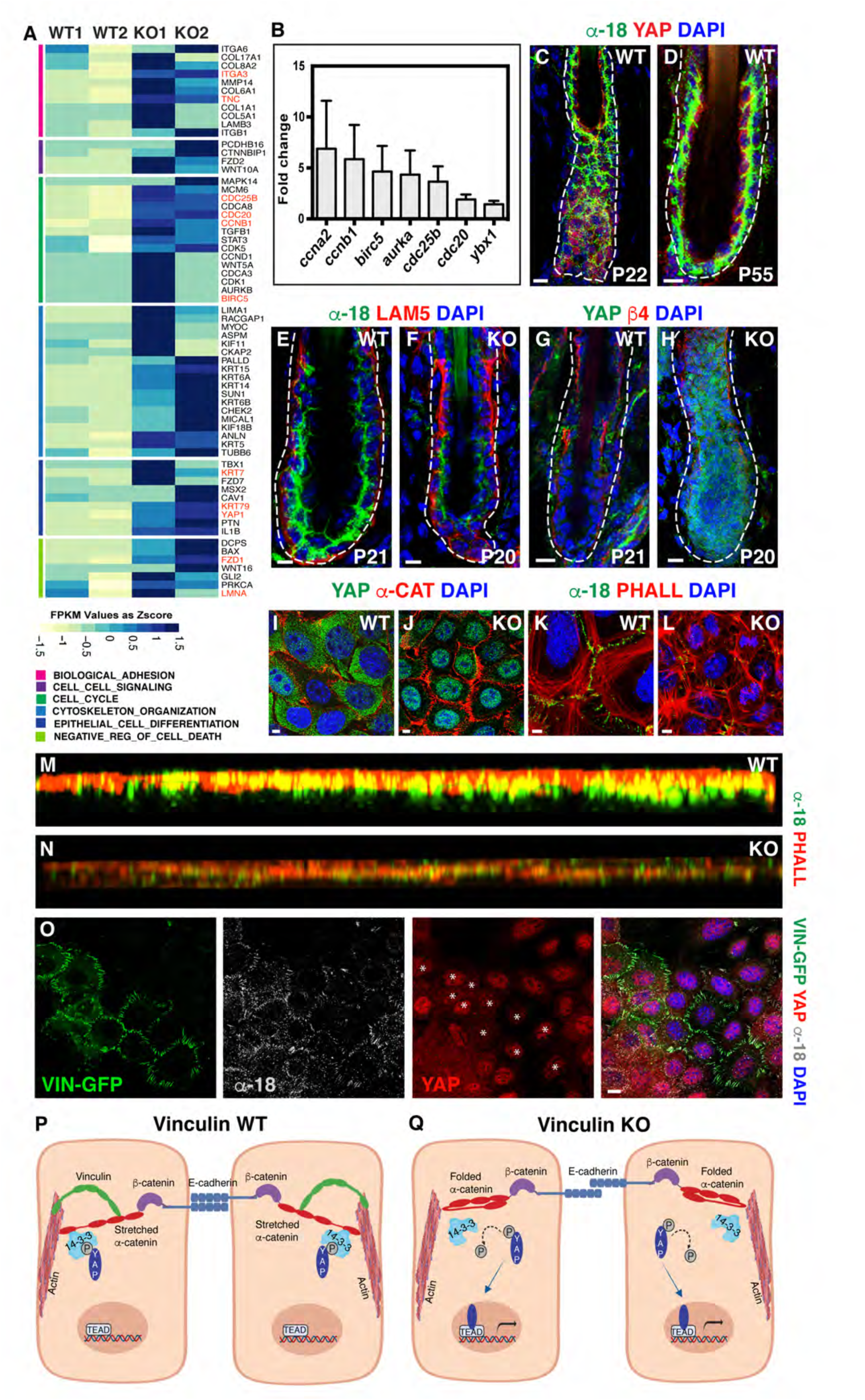
Loss of vinculin affects the conformation of **α**-catenin at AJs and disrupts its interaction with YAP. Heat map indicating the genes and pathways upregulated in vinculin KO bulge stem cells (A). Real time PCR analysis of Yap targets (B). WT (P-22) T/A transition and (P-55) telogen follicle junctions labeled with α-18 (green), YAP (red) and DAPI (blue) (C, D). WT and vinculin KO telogen follicles labeled with α-18 (green), lam5 (red) and DAPI (blue) in (E, F). Expression of YAP (green), β4-integrin (red) and DAPI (blue) in WT and vinculin KO telogen follicles (G, H). Expression of YAP (green), α-catenin and DAPI in confluent WT and vinculin KO cells (I, J). Expression of α-18 (green), phalloidin (red) and DAPI (blue) in confluent WT and vinculin KO cells (K, L). 3D SIM image of the expression of α-18 (green), phalloidin (red) at AJs (M, N). Vinculin KO keratinocytes transduced with vinculin-GFP, labeled with YAP1 (red), α-18 (grey) and DAPI (blue) (O) (asterisks represent the nuclei with reduced YAP1 expression). Model showing the relationship between the loss of junctional stability and increased signaling through nuclear Yap (P, Q). Scale bar: 10μM (C-H, O), 5μM (I-L).

YAP, a mechanosensor is a downstream effector of the Hippo pathway and shuttles to the nucleus to drive cellular proliferation (Low et al., 2014). Since YAP is a potent activator of the cell cycle, the upregulation of cell-cycle genes correlated with increased proliferation seen in the vinculin KO BuSCs. Upon contact inhibition YAP/TAZ translocates out of the nucleus and is maintained primarily at cell junctions (Zhao et al., 2007). Interestingly, α-catenin can sequester YAP at the junction, either through direct binding, (Sarpal et al., 2019; Silvis et al., 2011) or through its interaction with the adaptor protein 14-3-3 (Li et al., 2015; Schlegelmilch et al., 2011). Previous studies have reported that the loss of contact inhibition in α-catenin KO keratinocytes was associated with translocation of YAP into the nucleus (Silvis et al., 2011). Contact inhibited cells are associated with strong junctional expression of AJ proteins. Consistent with that notion, we observed a marked increase in the expression of α-catenin in the telogen follicles in comparison with the rapidly growing T/A follicles (Fig 1F, S1A). We also observed increased expression of the α-18 epitope antibody (which recognizes the stretched conformation of α-catenin) in the telogen follicles compared to the T/A follicles (Fig 6C, D) (Nagafuchi and Tsukita, 1994; Yonemura et al., 2010). Intriguingly this lower expression of α-18 in WT anagen follicles was associated with increased nuclear localization of YAP and increased proliferation (Fig 6C); whereas YAP expression remained predominantly cytoplasmic in the WT telogen follicles (Fig 6D). We investigated the expression of α-18 antibody in the bulge of WT and KO telogen follicles. The expression of α-18 was significantly reduced in the KO bulge compared to the WT (Fig 6E, F and Fig S6A, B). We then looked for the expression of YAP and its transcriptional co-activator TAZ in the BuSCs of WT and KO animals. We found increased nuclear expression of YAP and TAZ in the KO follicles, compared to a complete absence of nuclear signal in WT telogen follicles (Fig 6G, H, Fig S6C, D and E-H). Interestingly the total levels of YAP protein were not changed in the KO cells compared to the WT suggesting that the effects we see are probably due to a change in localization of the protein (Fig S6K). To correlate the loss of contact inhibition and the nuclear localization of YAP with the reduced expression of α-18 at the junctions, we cultured WT and vinculin KO keratinocytes to confluence. In KO keratinocytes, we observed strong nuclear localization of YAP, and increased, diffuse expression of α-catenin at the junctions compared to WT cells (Fig 6I, J). Interestingly these confluent KO cells displayed very weak expression of α-18, compared to the WT cells suggesting that the α-catenin at these junctions was not in a stretched conformation (Fig 6K, L). We also observed that the actin cytoskeletal network appeared disrupted in the KO cells, predominantly comprising of stress fibers (Fig 6L). We investigated the status of the connection of the stretched conformation of α-catenin with the actin cytoskeleton using structured illumination microscopy (3D SIM imaging). In the WT junctions α-18 colocalized with the actin filaments marked with phalloidin, indicating a strong association of α-catenin with the actin cytoskeletal network (Fig 6M). In the KO junctions however, there was minimal colocalization between α-18 and phalloidin, suggesting once again that in the KO junctions α-catenin is not maintained in a stretched conformation (Fig 6N). To confirm that the junctional localization of vinculin was critical to maintaining the stretched conformation of α-catenin and thereby sequestering YAP at the junctions, we transduced KO cells with either an empty Zs-Green vector, or the vin-GFP construct. In the mock-transduced cells, there was high expression of YAP in the nucleus (Fig S6 I). In the vin-GFP transduced cells on the other hand, we observed a significant decrease in the nuclear localization of YAP (white asterisks), and a concomitant increase in expression of α-18 at the junctions (Fig 6 O).

Taken together, we propose the following model. In WT contact inhibited cells, the binding of vinculin to α-catenin is essential to keep it in a stretched conformation (Seddiki et al., 2018; Yao et al., 2014), and associated with the actin cytoskeletal network. The stretched α-catenin creates a binding site for the adaptor protein 14-3-3, which in turn binds to p-YAP and sequesters it at the adherens junctions (Schlegelmilch et al., 2011) (Fig 6P). In vinculin KO cells on the other hand α-catenin is no longer maintained in a stretched conformation. This we believe disrupts the binding of α-catenin and 14-3-3, which is no longer able to sequester YAP at the AJs. YAP is dephosphorylated and is free to translocate into the nucleus and regulate cell cycle genes (among many of its targets) thereby affecting the loss of quiescence in vinculin KO BuSCs (Fig 6Q).

## Discussion

AJs have emerged as a mechanosensitive hub that have critical functions in regulating tissue development and maintaining homeostasis (Biggs et al., 2019; Fernandez-Sanchez et al., 2015; Miroshnikova et al., 2018; Pinheiro and Bellaïche, 2018; Vasquez and Martin, 2016). Mechanical forces applied on AJs are known to affect various important cellular functions such as stemness, proliferation, differentiation, collective cell migration, and apoptosis (Gilmour et al., 2017; Heisenberg and Bellaïche, 2013; Petridou et al., 2017). Thus, AJ remodeling under tension has been shown to be one of the key regulator of epithelial morphogenesis (Pinheiro and Bellaïche, 2018). Initially, E-cadherin and α-catenin were known to be the main regulators that contribute to the tension-dependent AJ reinforcement (Buckley et al., 2014; Perret et al., 2004; Rakshit et al., 2012). However, recently the idea of other AJ proteins being the modulators of AJs’ reinforcement under tension has started to emerge. In line with this idea, we have described a hitherto unknown mechanism of the role of vinculin in maintaining stem cell quiescence.

Based on *in vitro* experiments, vinculin has been shown to be a potent mechano-transducer, however this has not seemed to be the case *in vivo* (Goldmann, 2016; Martino et al., 2018; Spanjaard and de Rooij, 2013). While the *in vitro* manipulation of vinculin at FAs have profound consequences on the forces generated by these junctions, the *in vivo* function of vinculin has remained unclear because its genetic loss-of-function in worms, flies and fish elicit weak or no phenotypes (Alatortsev et al., 1997; Barstead and Waterston, 1989; Han et al., 2017; Xu et al., 1998). In mice, the loss of vinculin results primarily in cardiac defects (Xu et al., 1998). The heart is an organ that undergoes repeated contraction, and therefore may be more susceptible to changes in the mechanical strength of its junctions (Lyon et al., 2015; Sheikh et al., 2009). Interestingly, when we conditionally deleted vinculin from the epidermis using the K14-Cre driver, we noticed a striking effect on hair cycling, which also is a phenomenon associated with repeated mechanical stress associated with the growth, regression and the resting phase of the hair follicle. Specifically, we observed a distinct loss of the telogen or resting phase (starting form the second hair cycle) and perturbation in hair follicle cycling starting from postnatal day 10, resulting in the eventual thinning of the hair coat. To better understand the hair cycling phenotypes in the vinculin KO animals, we performed a detailed and comprehensive analysis of hair cycling between P1-P64. Our study allowed us to discern that the HF cycle in vinculin KO skin was markedly accelerated, such that the KO animals complete two hair cycles by P37, whereas the second hair cycle in the WT ends at P110. To the best of our knowledge, this is the first animal model, where the hair follicle cycling is so dramatically accelerated. Intriguingly, our analysis of the α-catenin KO skin suggests that the HF cycle was also accelerated in these animals, albeit with some differences. The telogen stage in the α-catenin KO occurs at P27, compared to P21 in the WT and the follicles in general appear to be smaller than those in the vinculin KO. The difference in the hair cycle between the α-catenin and vinculin animals could be attributed to the cre driver line, and the efficiency with which the expression α-catenin and vinculin are lost. Interestingly, in both the vinculin and α-catenin KO animals; there was a loss of the CD34, LRIG1-positive quiescent pools of stem cells, the ectopic expression/expansion of stem or progenitor cell markers such as SOX9 and K15, and an accelerated entry into the cell cycle as evident by increased Ki67 staining and other cell cycle markers. Taken together these results suggest a critical role for AJ proteins in the maintenance BuSC quiescence.

### Junctional strength of vinculin and α-catenin KO cells is similar

The forces required to detach an E-cadherin coated bead from vinculin KO and α-catenin KO cells was ∼2.5nN compared to ∼5-6nN in WT cells. How does one explain this observation? Reinforcement of the junctions can occur through the formation of catch bonds between cadherin molecules and between α-catenin and actin and vinculin and actin in a force dependent manner. In the α-catenin KO, although E-cadherin and β-catenin can localize to the membrane, there is no recruitment of vinculin to the AJs. In such a scenario, the only reinforcement at junctions might occur by E-cadherin homotypic interactions, without reinforcement through the actin machinery, since both proteins that can contribute to the catch bond formation are missing (Buckley et al., 2014; Perret et al., 2004; Rakshit et al., 2012; Swaminathan et al., 2017). In vinculin KO cells on the other hand, there is increased expression of α-catenin at the junctions, possibly as a compensatory mechanism, however this α-catenin does not appear to be in a stretched conformation. This was seen by labeling the junctions in the vinculin KOs with the α-18 antibody both *in vivo* and *in vitro*. In both cases, there was a severe reduction in the level of α-18 at the AJs, suggesting that the loss of vinculin resulted in the failure of α-catenin to be maintained in the stretched conformation. This in turn could interfere with the ability of α-catenin to form a catch bond with the actin network, thus weakening the junctions. The force of 2.5nN required to dislodge the E-cadherin coated beads from both the vinculin and α-catenin KO cells probably represents the strength of E-cadherin mediated catch bonds that remain unchanged in both KO cells.

### Junctional stability and bulge stem cell quiescence

One of our confounding results was the restriction of the phenotype of the vinculin KOs to the BuSC compartment with little or no effect seen in the IFE. This made us wonder about the nature of the stem cell niche itself, particularly with respect to the AJs. While the maintenance of the niche has been ascribed to multiple cell-intrinsic factors such as SOX9, LGR5, K15 etc, the loss of AJ proteins from the skin seem to have a pleotropic effect. However, when we carefully examined the expression of AJ proteins in the bulge, we noticed a significant increase in their expression, associated with a strong cortical actin meshwork. This expression was reminiscent of the staining of AJ proteins and the actin network in contact inhibited cells *in vitro.* This contact inhibited state, in turn, keeps YAP, a potent activator of the cell cycle, in the cytoplasm, by sequestering it at the junctions. Intriguingly, NGS analysis of transcripts expressed in the vinculin KO bulge cells revealed that both YAP1 and its targets, the cyclins, were upregulated in the KO. Analysis of the KO telogen follicles further revealed that increased proliferation seen in the KOs was associated with the nuclear expression of YAP1. The localization of YAP1 to the junctions occurs though its interaction with α-catenin (Schlegelmilch et al., 2011). In the vinculin KO bulge cells, although there was increased expression of α -catenin at the AJs (possibly as a compensatory mechanism), YAP failed to be sequestered to these junction, due to the fact that α-catenin was no longer in a stretched conformation. This suggests that AJ stability, an important prerequisite to establish contact inhibition, relies not just on the presence of all of the component proteins, but also on the conformational states of these proteins. The loss of contact inhibition in the vinculin KO bulge, results in failure to acquire a quiescent state and the cells of the bulge compartment proliferate in a manner similar to the cells of the growing anagen follicles. In fact the vinculin KO follicles seem to cycle between anagen and catagen stages, completely bypassing the telogen stage, or establishing a club hair. Based on these findings, we propose that one of the mechanisms to maintain this quiescent state is through contact inhibition.

The maintenance of this contact inhibited state therefore may need to be balanced with the requirement of cells in the hair follicle to proliferate rapidly at the start of a new hair cycle. One way that this balance can be maintained is through cadherin switching. In the secondary hair germ, the first part of the follicle that grows in the new hair cycle, the dominant cadherin is P-cadherin (Nanba et al., 2000). The junctions nucleated though this cadherin have lower junctional stability associated with higher proliferative potential. As the follicles grow, the junctions become more labile, and this in turn allows YAP to translocate into the nucleus providing the proliferative burst needed for growth of the HF. Interestingly even as the HF is rapidly growing, the bulge compartment maintains stable E-cadherin mediated junctions (Rognoni and Walko, 2019).

In the proliferative basal layer, the AJs are more labile and associated with lower levels of E-cadherin, and as such these cells are not contact inhibited. The loss of vinculin in the basal keratinocytes may not be as critical and therefore may not result in any discernable affects/phenotypes. The suprabasal cells that are contact inhibited also do not seem to be affected in the vinculin KOs. This could be due to two reason, firstly, the suprabasal cells are post-mitotic, and do not have significant expression of YAP. Therefore instability of these junctions would not result in nuclear localization of YAP. Secondly the number of desmosomes in the KO IFE does not vary significantly from that seen in the WT IFE (as opposed to the significant decrease of desmosomes in KO HFs). These desmosomes may provide additional junctional stability to the suprabasal cells, thus mitigating the loss of vinculin in the IFE.

Taken together, our data clearly implicate the role of junctional stability and contact inhibition in maintaining BuSC quiescence and suggests that this may override and/or regulate other cell intrinsic mechanisms that are in place to keep the stem cells in a quiescent state. Given the importance of stem cell re-activation in cancer, the results from this study may have critical implications in the regulation of cell-cell adhesion, contact inhibition and stemness during skin tumorigenesis, and tumor progression.

## Materials and Methods

### Mice

Vinculin floxed mice were obtained from the Ross lab at UCSD (Zemljic-Harpf et al., 2007) and crossed with K14 Cre mouse in C57BL6 background. Animals were bred in the NCBS/inStem Animal Care Resource Centre and the all procedures were approved by the inStem Institutional Animal Ethics Committee.

Mice were injected intra-peritoneally at P4 and P5 with 25ug/gm weight of 5-Ethynyl-2-deoxyuridine (EdU) (Thermo Fisher Scientific) and chased for times specified (Vin WT P-21 and P-53 and Vin KO P-20 and P-36). To detect the EdU, click-iT EdU Flour 488 imaging kit was used.

### Cell Culture

WT, Vinculin and α-catenin KO keratinocytes were isolated as described in (Raghavan et al., 2003) and were grown in high calcium E media (DMEM/F12 in a 3:1 ratio with 15% FBS supplemented with insulin, transferrin, hydrocortisone, cholera toxin, triiodothyronone and penicillin/streptomycin at 32°C.

### Hair and Skin Reconstitution assay

Hair and skin reconstitution assays were performed as described in (Blanpain et al., 2004). Briefly, 8-week-old male NSG mice were used as recipients. A wound of ∼ 1.5 cm diameter was made on the back of anesthetized NSG mice, and a sterile silicon chamber was inserted. A slurry of 100 −200 μl of cell suspension containing freshly isolated or cultured epidermal keratinocytes with freshly isolated or cultured dermal fibroblasts were added into the chamber through the top hole of the chamber. Approximately 4 million keratinocytes and 8 million fibroblasts were added to make the slurry. The chamber was removed after 1 week and the wounded or graft site was harvested 5 to 6 weeks after transplantation.

### Lentiviral transduction of keratinocytes

All the vinculin rescue constructs were cloned into the pLVV-IRES Zs-Green lentiviral vector (Clontech). The Lentiviruses were generated through transient co-transfection of the vector plasmid, the packaging construct pCMVD8.9 and the envelope coding plasmid pCMV-VSVG (from Dr. Soosan Ghazizadeh, SUNY Stony Brook) into 293T cells using lipofectamine 2000 (Invitrogen). Supernatant containing the virus was collected after 72 hours and filtered through a 0.22 micron PES (Millipore) filter. The MOI was calculated by infecting 293T cells and ranged from 5×10^5^ - 1x 10^6^ IU. Vinculin knockout keratinocytes were plated on the previous day at a low density (15-20% confluent). The media was removed from the KO keratinocytes and the filtered viral supernatant containing 8 μg/ml polybrene was added and incubated for 5 hours at 32°C. After 5 hours the virus was removed and fresh E media added.

### Hair cycle analysis and shaving Experiment

The detailed hair cycle analysis was done based on (Müller-Röver et al., 2001). Tissues were fixed with Bouin’s fixative and embedded in paraffin and sectioned. Sections were stained with hematoxylin/eosin and imaged using an IX73 Olympus microscope. For shaving experiments, P53 WT and P36 vinculin KO animals were used. The dorsal pellage hair of the animals was shaved under anesthesia and the regrowth of hair was observed and photographed after one week.

### Immunofluorescence and Immunohistochemistry

Approximately 5-10,000 cells were plated on glass coverslips coated with 10μg/ml fibronectin for 24 hours. Cells were fixed with 4% paraformaldehyde for 10 minutes at room temperature (RT), permeabilized with PBST (PBS+ 0.2% triton-x-100) for 10 minutes and blocked with 5% normal donkey serum in PBST for 1 hour at RT. The cells were incubated with the primary antibody for 1 hour at RT, washed 5 times with PBS and incubated with Alexa fluor 488, 568 and 647 secondary antibodies (1:250; Invitrogen) for 30 minutes at RT. The nuclei were counterstained with 4’, 6 -diamidino-2-phenylindole (DAPI). For the skin sections, the primary antibody was incubated overnight at 4°C, followed by 1h secondary antibody incubation. Antibodies and dilution used were: - Vinculin (1:100), α-catenin (1:1000), β-catenin (1:1000) (Sigma), K14 (1:1000), K6 (1:100), Involucrin (1:100), (gift from Satrajit Sinha), E-cadherin (1:1000), Phalloidin 488, 568, 647 (1:100), (Thermo Fisher Scientific), P-cadherin (1:200, Invitrogen), K5 (1:200, Biolegend), Laminin 5 (1:1000, gift from Bob Burgeson), α-SMA (1:100), K15 (1:200), Lgr5 (1:200), Ki67 (1:200), AE13 (1:100), AE15 (1:100), YAP (1:100) (Abcam), Sox-9 (1:200, Millipore), Integrin Beta 4 (1:100, BD), Lrig1(1:500, R&D System), Lef1 (1:200), YAP1 (1:100) (Cell Signaling), TAZ (1:100, gift from Marius Sudol)

### Flow analysis cell sorting

Stem cells were isolated from P21 WT and P20 KO telogen follicles as described in (Tumbar et al., 2004). Cells were labeled with CD34-FITC and α6 integrin-APC conjugated antibodies, isotype control antibodies (eBioscience) and propidium iodide (Sigma). The cells were sorted using the BD FACS Aria Fusion sorter. Total RNA was extracted from WT and vinculin KO cells directly collected in Trizol LS (Invitrogen). RNA quality was determined by using a Bioanalyzer.

### RNA Sequencing and qRT-PCR

Single-end RNA sequencing (1×75bp) was performed on pooled WT and vinculin KO bulge stem cell samples. Nextseq500 was used for sequencing from the cDNA libraries. ∼31 to 40 million reads were obtained per sample. Trimmomatic adapters were used (Bolger et al., 2014) for mapping them to rRNA. Reads that did not align to rRNA were taken for further analysis. Reference based transcriptome assembly algorithms Hisat2 v2.1.0 (Kim et al., 2015); Cufflinks v2.2.1 (Trapnell et al., 2010) and Cuffdiff v1.3.0 (Trapnell et al., 2013) were used to identify differentially expressed transcripts. The reads were aligned to mouse (mm10) genome-using Hisat2 (-q -p 8 --min-intronlen 50 -- max-intronlen 250000 --dta-cufflinks --new-summary --summary-file). Around, 50-60% of reads mapped back to reference genome. The mapped reads were assembled using Cufflinks using mm10 Refseq gtf file. Since every replicate sequenced was a pool from multiple samples (animals) and considering intrinsic variation of KO between animals, differential expression was done without pooling the replicates using Cuffdiff v1.3.0. Genes which had adjusted p-value < 0.05 & minimum two-fold up/down regulated were considered as significantly expressed and taken for further analysis. To increase the stringency, FPKM filter were used to remove low expressed transcripts. Significantly expressed genes were overlapped from individual replicate comparison (WT1 Vs. KO1, WT1 Vs. KO2, WT2 Vs. KO2, WT2 Vs. KO1) and identified 2002 transcripts, which were significantly expressed in at least two comparisons (Supplementary table 1,2). 878 transcripts identified that were two-fold up-regulated and 1124 transcripts were two-fold down-regulated between Vin WT and Vin KO animals.

Pathway analysis & gene-ontology analysis were done for these selected up/down regulated transcripts using GSEA (Mootha et al., 2003; Subramanian et al., 2005). Customized perl script was used for all the analysis in this study. Rggplot2 (Wickham, 2016), pHeatmap (https://cran.r-project.org/web/packages/pheatmap/index.html) & CummeRbund (Goff et al., 2013) library were used for plotting.

1 µg of total RNA was used to synthesize cDNA using SuperScript III RT First Strand cDNA Synthesis Kit (Invitrogen). Real-time PCRs were performed on an ABI7900HT system. mRNA expression level was quantified by using the ΔΔCt method and the gene expression was normalized to GAPDH. The list of primers used is provided in Supplementary Table 4.

### Immunoblotting

Whole skin tissue from WT and KO animals were snap-frozen in liquid nitrogen and pulverized using a tissue smasher. Cultured keratinocytes from WT and KO were washed with PBS after aspirating media. Cells were scraped using a rubber spatula. Both tissue and cells were re-suspended in RIPA lysis buffer (Sigma-Aldrich) containing protease inhibitor cocktail (Roche) and incubated on ice for 10 mins. The lysates were then centrifuged at high speed for 15 min and the supernatant was collected. The protein concentration was estimated using a BCA kit (Promega). The blots were blocked using 5% NFDM (Santa Cruz) and incubated with primary antibody overnight at 4°C. The blots were washed with 1× TBST and incubated with HRP conjugated secondary antibodies (1:3000, CST) for 1h at RT. The blots were developed using chemiluminescence in ImageQuant LAS 4000 mini or ibright biomolecular imager. Band intensities from the blots were quantified using Image J.

The following antibody concentrations were used for immunoblotting: -

α−catenin (1:1000), β−catenin (1:2500), Vinculin, E-cadherin (1:800), YAP1 (1:1000) (SantaCruz), Tubulin (1:1000), (Sigma), HRP anti-Rat Secondary (1:3000), HRP anti-Mouse Secondary (1:3000), HRP anti-Rabbit Secondary (1:3000), (CST).

### Ultrastructure Analysis

Back skin from anterior region of mice were fixed in 2% glutaraldehyde, 4% PFA, and 2 mM CaCl_2_ in 0.05 M sodium cacodylate buffer, pH 7.2, overnight at 4°C, post-fixed in 1% osmium tetroxide (OsO4). Skin sections were then counterstained with 4% uranyl acetate in water and processed for Epon embedding. Ultrathin sections (80-100 nm) were taken using RMC Cryo UltraMicrotome. For scanning EM of cultured keratinocytes on coverslips, cells were fixed with 2% glutaraldhehyde and 2mM CaCl_2_ (in 0.08M sodium-cacodylate pH 7.2) at room temperature for 20 mins post-fixed with 0.8% K3Fe(CN)6 and 1% OsO4 in 0.1M sodium-cacodylate for 30 mins on ice followed by block staining with 4% uranyl acetate in water. Cells were then serially dehydrated with ethanol and propylene oxide. After dehydration samples were dried using critical point-drying and coated with gold using sputter coating. Electron micrographs were taken in MERLIN Compact VP scanning electron microscope.

### Force measurements at the AJs

Briefly, 3μm microbeads coated with E-cadherin were scattered (using a 200ul microtip) on the fibronectin-coated coverslip where cells were grown. Individual microbeads were captured with an elastic glass micropipette and allowed to contact the cell surface for 5 minutes to allow AJ formation. The force (F) at which the bead was detached from the cell surface was a direct readout of the strength of the E-cadherin mediated AJs formed by these cells with the beads. The detaching force was measured and presented as shown in Figure 5 (Movie M5 - M11). This helped us to quantify the contribution of vinculin and α-catenin to the adhesion strength at the AJs. Since the strength of the AJs may depend on the contact time and the location on the cell surface, we chose a fixed 5-minute contact time and similar contact points that are located ∼5um away from nucleus on each cell.

### Live Cell Imaging

For live cell imaging, cells were cultured on glass bottom dish coated with fibronectin (10μg/ml). Imaging was performed on a motorized Nikon Eclipse Ti-E inverted microscope equipped with a Plan Fluor 40x/1.3 oil immersion objective lens, a 491nm laser (FRAP-3D laser launch; Roper, France), CSU-22 scanning head (Yokogawa; Emission filter 505-545), Evolve EM-CCD camera (Photometrics) and the Nikon Perfect Focus System (PFS). The cells were maintained at 37^°^C in a 5% CO2 humidified environment using an on-stage incubator (Live Cell Instrument, Chamlide). Both confocal and brightfield datasets were acquired at each time point. Time-lapse imaging was controlled through MetaMorph software (Molecular Devices). The image visualisation and analysis was completed using the Fiji (ImageJ) (Ferreira and Rasband, 2012; Schindelin et al., 2012) software package and Imaris (Oxford Instruments).

### Statistical Analysis

GraphPad v6.0e (Graphpad Software) was used to perform all the statistical analyses. Two tailed unpaired t test was used for comparing two groups of data sets. Mean, SD, SEM values were used to plot error bars.

## Supporting information

NGS_Analysis

Upregulated and Downregulated Pathways

Gene Ontology Analysis

## Author Contributions

SR designed the study. SR, RB, AB and YJ designed the experiments. SR, RB, AB, SL, ZZ, MN and VK performed the experiments. VL and DP analyzed the NGS data. GW& SR performed the live cell imaging. YJ and ZZ developed the adhesion force measurement assay and analyzed the force data. SR, YJ, DP, VV, RB, AB, SL, ZZ, MN, VL and VK analyzed the data. RB, AB, YJ, VV and SR wrote the manuscript.

## Acknowledgements

We would like to thank members of the Raghavan Lab, Ramanuj DasGupta, and Doina Tumbar for providing critical feedback on the work and manuscript. We thank H. Krishnamurthy and the Central Imaging and Flow Facility (CIFF) at NCBS/inStem for the use of the confocal microscopes, EM and FACs facility. We thank Awadesh Pandit at the NCBS NGS facility for performing NGS. We would like to give a very special thanks to Dr. Sangeetha at the NCBS animal facility for help with the hair and skin reconstitution experiments. We thank Prof. Satyajit Mayor (National Center of Biological Science (NCBS), India) for the vinculin constructs, Prof. Marius Sudol and Dr. Tony Kanchanawong (Mechanobiology Institute, NUS, Singapore) for the Yap antibodies and α-catenin constructs respectively and Dr. Akira Nagafuchi (Nara Medical University, Japan) for the α-18 antibody This work is supported by grants from the Science & Engineering Research Board (SERB) EMR/2016/003199, Department of Science and Technology (DST), India, and Institute for Stem Cell Biology and Regenerative Medicine (InStem), India core funding to SR. The work at Singapore is funded by the National Research Foundation, Prime Minister’s Office, Singapore, under its NRF Investigatorship Programme (NRF Investigatorship award no. NRF-NRFI2016-03) and the National Research Foundation, Prime Minister’s Office, Singapore and the Ministry of Education under the Research Centres of Excellence Programme (to J.Y.). RB is supported by a DST-SERB WOS-A predoctoral fellowship (SR/WOS-A/LS-255/2017). AB was funded through a NPDF postdoctoral funding from DST SERB (PDF/2017/000860). Animal work was partially supported by the National Mouse Research Resource (NaMoR) grant (BT/PR5981/MED/31/181/2012; 2013-2016) from the DBT.

---

**Figure S1:**
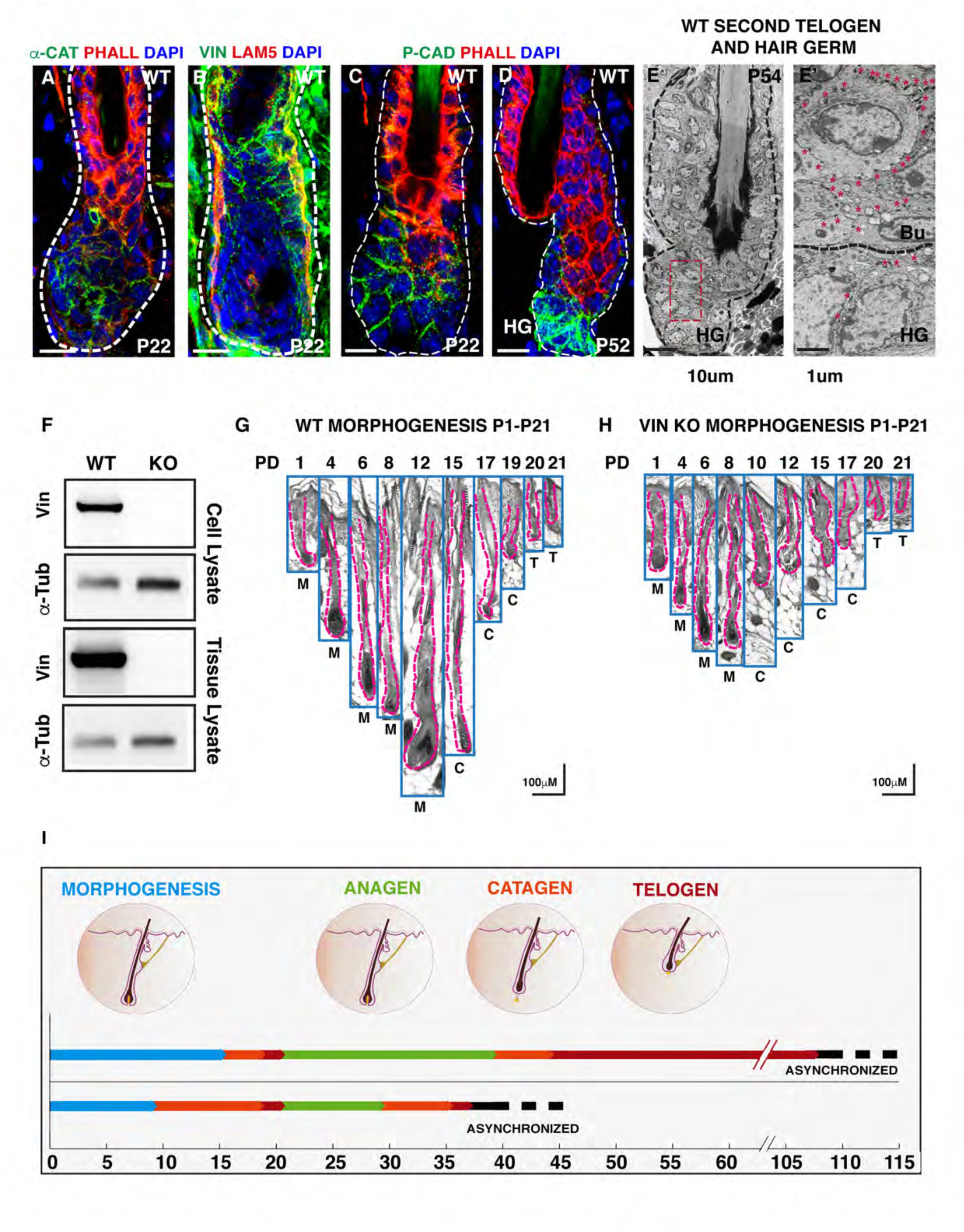
Analysis of the hair follicle cycle. WT (P-22) T/A transition follicle junctions labeled with α-catenin (green), phalloidin (red) and DAPI (blue) (A) and vinculin (green) and laminin 5 (red) DAPI (blue) (B). WT Telogen-Anagen (T/A) transition (P-22) and Telogen (P-52) follicles labeled with P-cadherin (green), phalloidin (red) DAPI (blue) (C, D). SEM of the cell-cell junctions at telogen (WT, P-54) follicles (E). Insets: high mag images of the areas indicated by the red dashed box (E’). Red asterisks indicate desmosomes. HG-Hair germ, Bu-Bulge. Immunoblot analysis of vinculin protein expression from WT and vinculin KO keratinocyte and epidermal lysates (F). Histological analysis of the hair follicles from WT and vinculin KO animals from morphogenesis to the telogen phase (G, H). Schematic representing the first two hair cycles in WT and vinculin KO animals (I). Scale bar: 10μM (A-E), 1μM (E’), 100μM (G, H).

**Figure S2:**
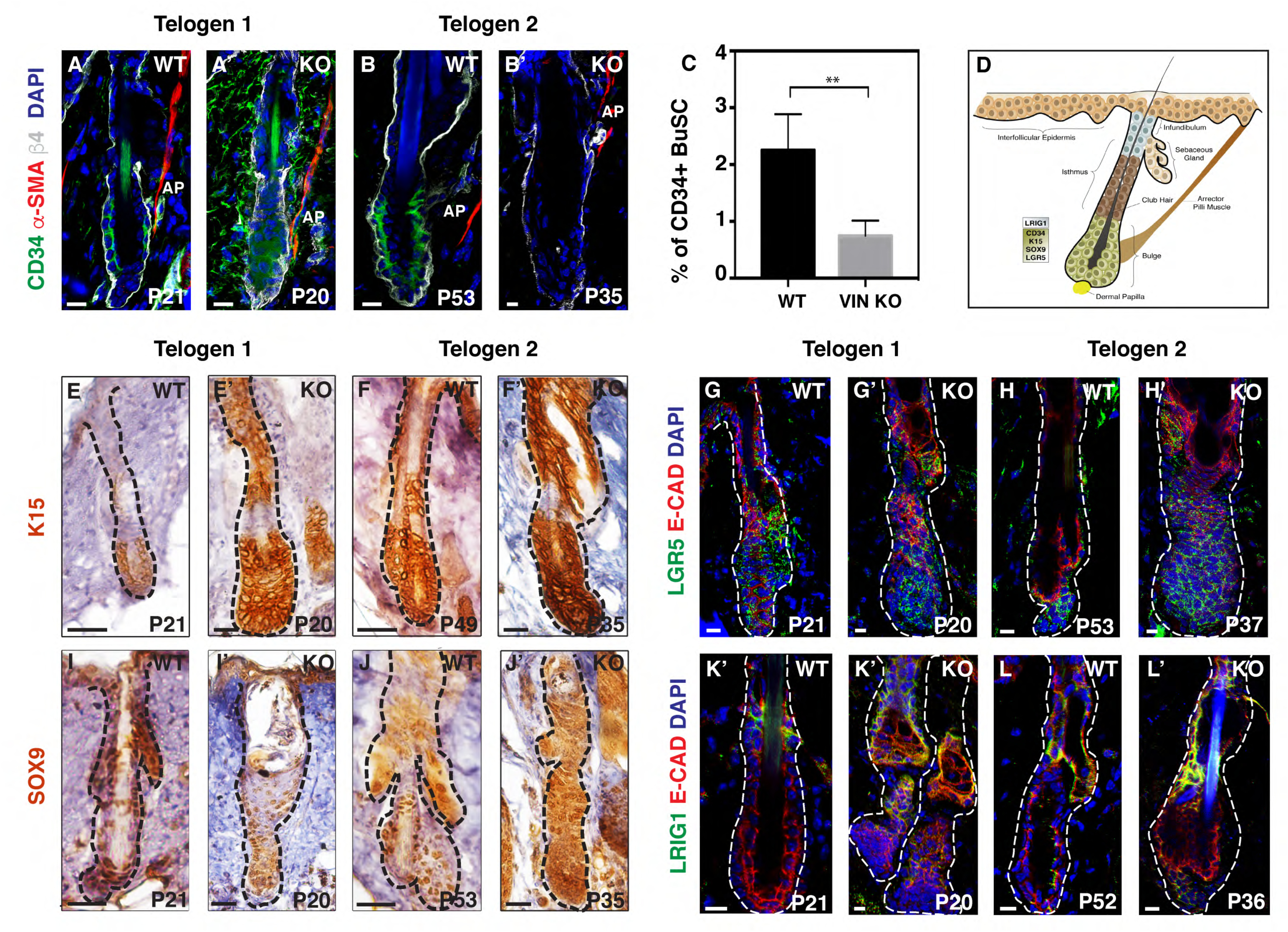
Analysis of the bulge stem cell compartment in first and second telogen follicles: WT and vinculin KO bulge stem cells labeled with CD34 (green), a-SMA (red), b4 integrin (grey) and DAPI (blue) (A, A’, B, B’). Histogram representing the percentage of BuSCs by FACS in WT and Vin KO epidermis (C). Schematic of the bulge compartment indicating location of the different stem cell pools (D). Immunohistochemistry of WT and vinculin KO bulge stem cells labeled with K15 (E, E’, F, F’). Immunohistochemistry of WT and vinculin KO bulge stem cells labeled with Sox9 (I, I’, J, J’). WT and vinculin KO bulge stem cells labeled with Lgr5 (green), E-cad (red) and DAPI (blue) (G, G’, H, H’). WT and vinculin KO stem cells labeled with Lrig1 (green) E-cad (red) and DAPI (blue) (K, K’, L, L’). Scale bar: 10 μM (A-B’, G-L’), 20 μM (E-J’). \*\**P* < 0.01.

**Figure S3:**
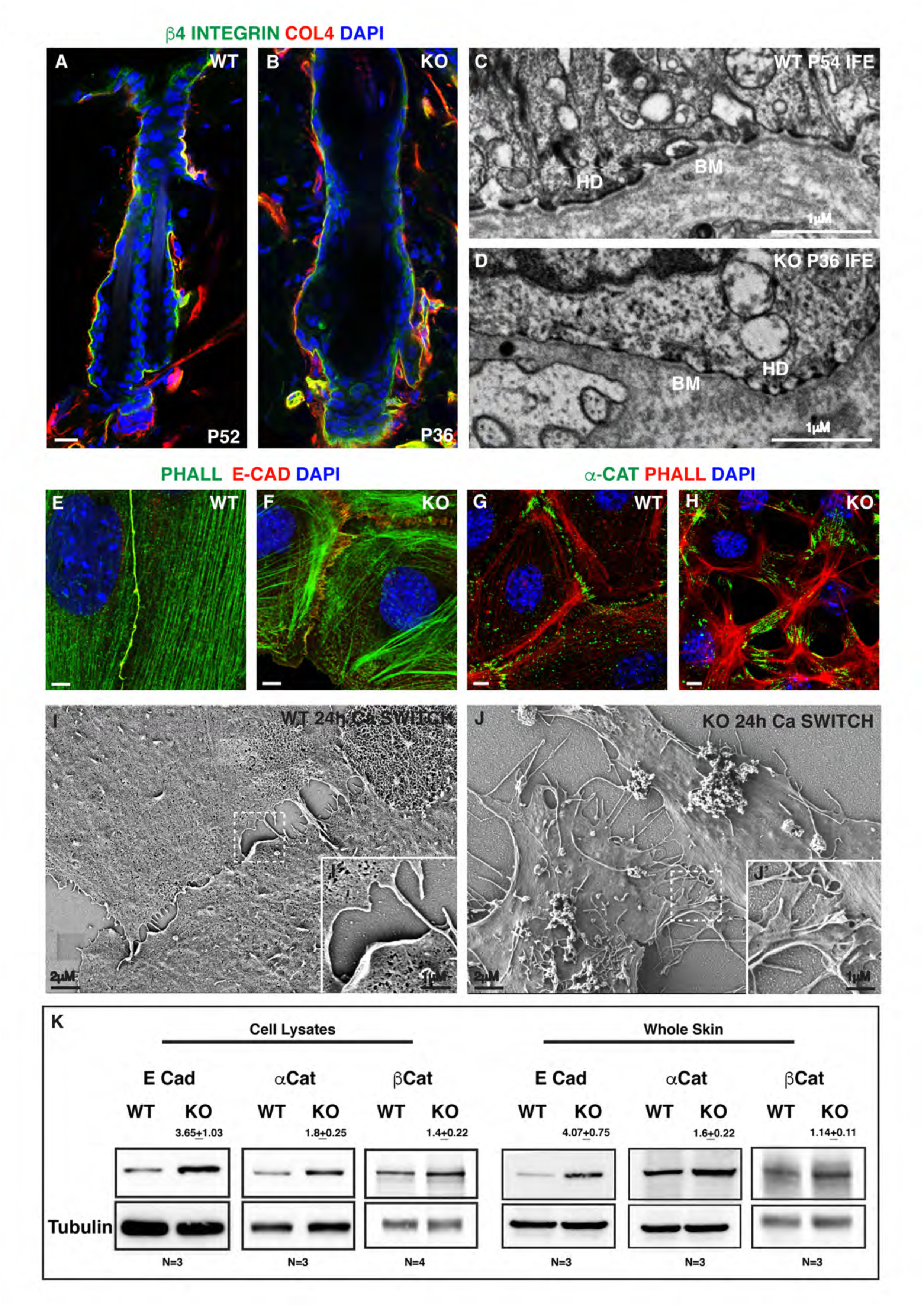
Status of the junctions in vinculin KO telogen follicles and keratinocytes. Expression of β4-integrin (green), collagen IV (red) and DAPI (blue) in WT and vinculin KO telogen follicles (A, B). Electron micrographs of the basement membrane in the IFE of WT and vinculin KO telogen follicles (C, D). Adherens junction formation after 24hrs of calcium switch in WT and vinculin KO keratinocytes marked with E cadherin (red), phalloidin (green), DAPI (blue) (E, F). Junction formation in WT and vinculin KO keratinocytes marked with α-catenin (green), phalloidin (red) and DAPI (blue) (G, H). Scanning electron micrographs of junction formation after 24 hours of calcium switches in WT and vinculin KO keratinocytes (I, J). Insets: high mag images of the junctions indicated by the dashed box (I’, J’). Immunoblot analysis of adherens junction proteins from WT and vinculin KO keratinocyte and whole skin lysate (K). HD-hemi desmosome, BM-basement membrane. Scale bar: 10μM (A, B), 1μM (C, D), 5μM (E-H), 2μM (I, J), 1μM (I’, J’).

**Figure S4:**
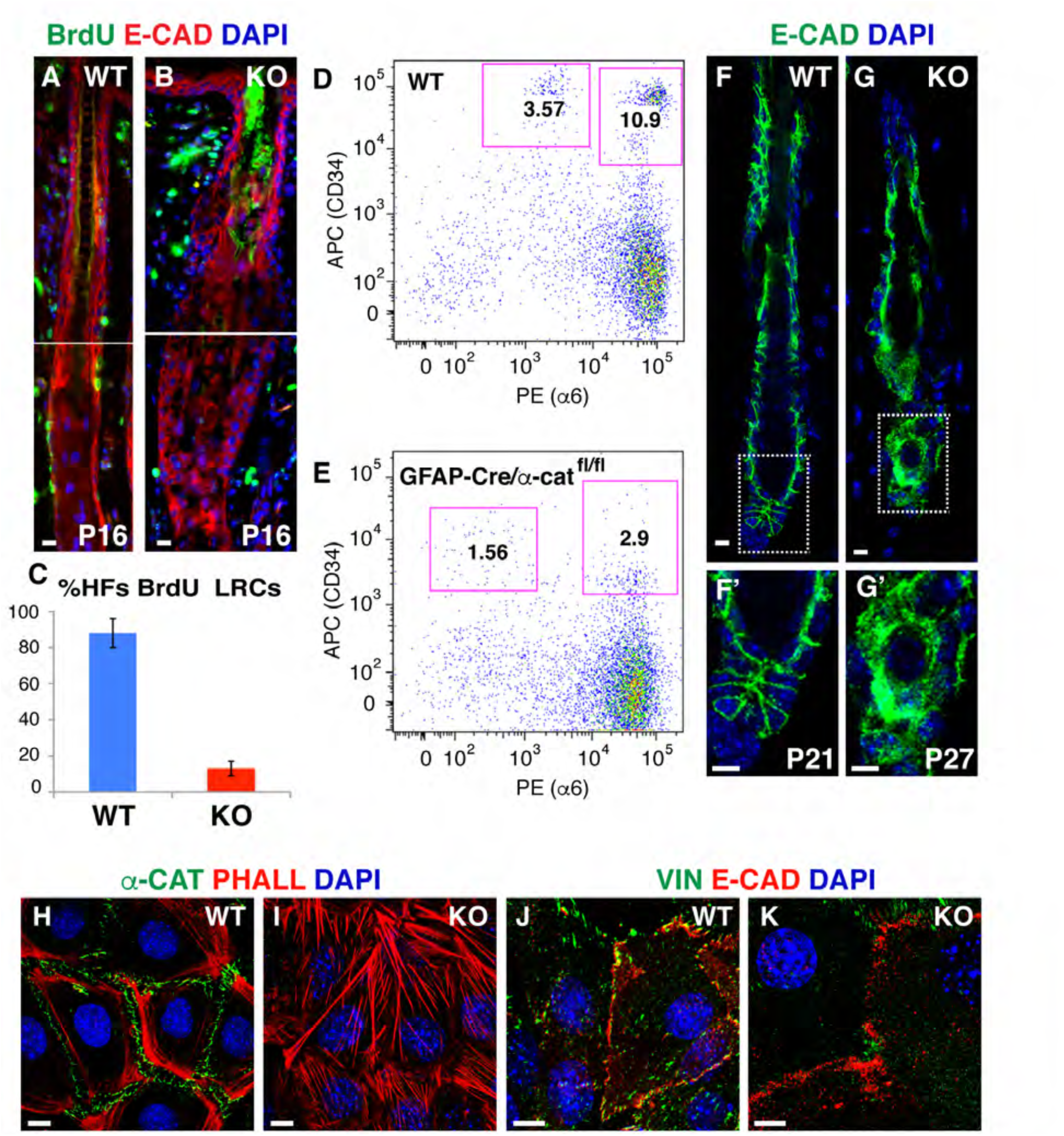
Analysis of stem cells and AJs in α-catenin KO. WT P-16 and α-catenin KO P-16 bulge stem cells labeled with BrdU (green), E-cadherin (red) and DAPI (blue) (A, B). Histogram representing the number of BrdU positive LRCs in WT and α-catenin KO bulge compartment (C). FACS analysis of bulge stem cell population in WT and α-catenin KO (D, E). Back skin sections from 7-8.5M WT and α-catenin KO animals labeled with CD34 (green), E-cadherin (red) and DAPI (blue) (D, E). AJ in WT and α-catenin KO telogen follicles labeled with E-cadherin (green) and DAPI (blue) (F, G). High mag images of the junctions indicated by the dashed box (F’, G’). WT and α-catenin KO keratinocytes labeled with α-catenin (green), Phalloidin (red) and DAPI (blue) (H, I). Junctions in WT and α-catenin KO keratinocytes marked with Vinculin (green), E-cadherin (red) and DAPI (blue) (J, K). Scale bar: 5μM (A, B, F, G, F’, G’), 10 μM (H-K).

**Figure S6:**
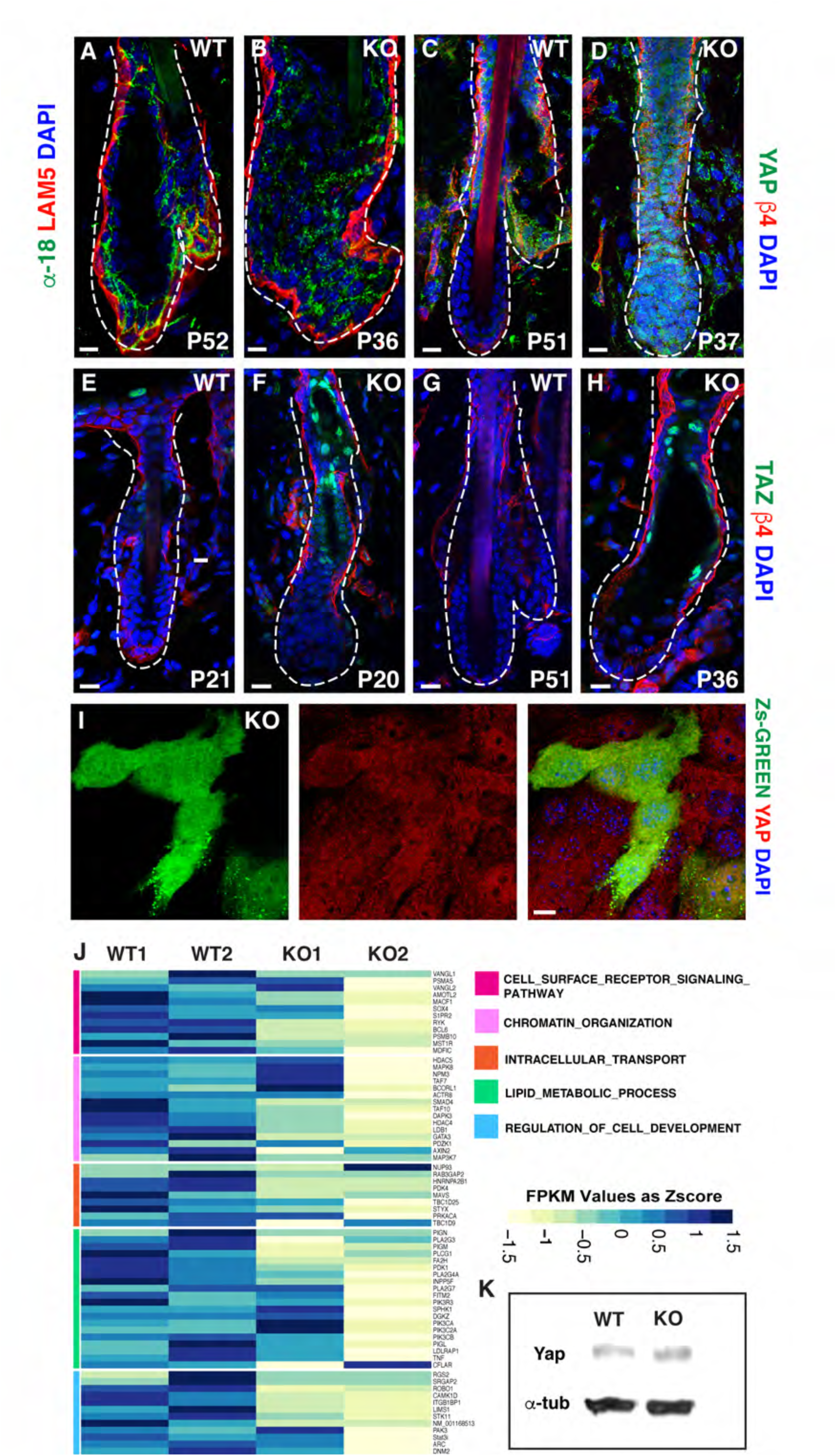
Expression of YAP and TAZ in WT and vinculin KO. Expression of α-18 (green), Laminin 5 (red) and DAPI (blue) in WT and vinculin KO 2^nd^ telogen follicles (A, B). Expression of Yap (green), β4-integrin (red) and DAPI (blue) in WT and vinculin KO 2^nd^ telogen follicles (C, D). Expression of TAZ (green), β4-integrin (red) and DAPI (blue) in WT and vinculin KO 1^st^ and 2^nd^ telogen follicles (E-H). Vinculin KO keratinocytes transduced with Zs-green control vector labeled with Yap (red) and DAPI (blue) (I). Heat map of pathways down regulated in Vinculin KO bulge stem cells transcriptome (J). Immunoblot analysis of YAP expression in WT and vinculin KO keratinocytes (K). Scale bar: 10 μM (A-I).

**Extended supplementary Figure ES1:**
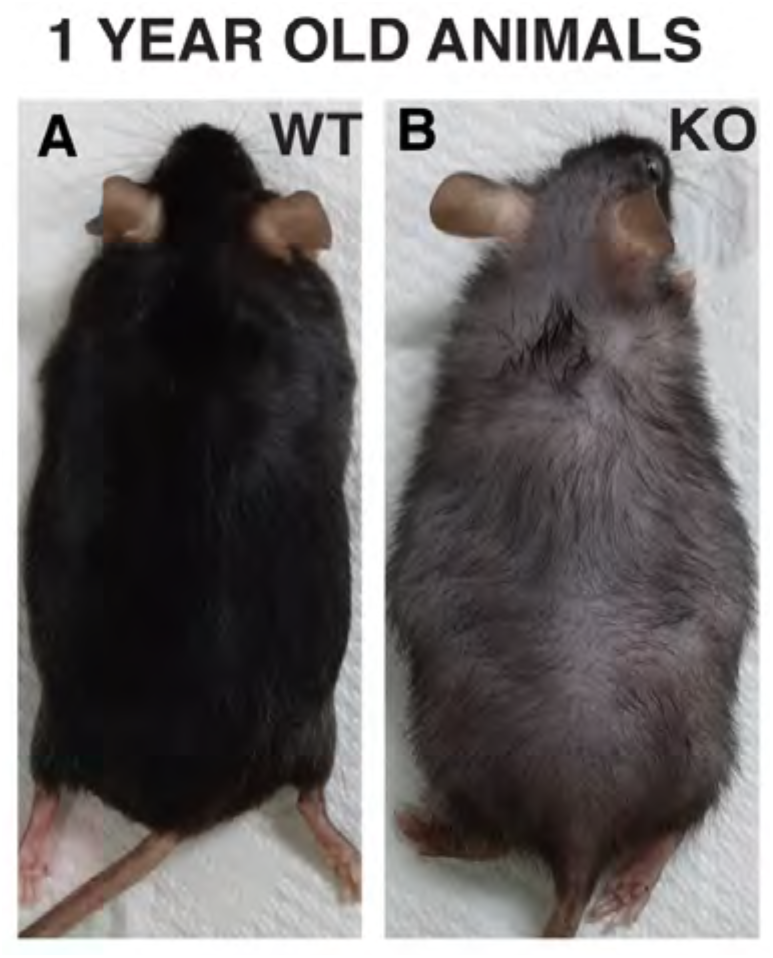
Phenotype of 1-year old WT and vinculin conditional KO animals (A, B).

**Extended supplementary Fig 2:**
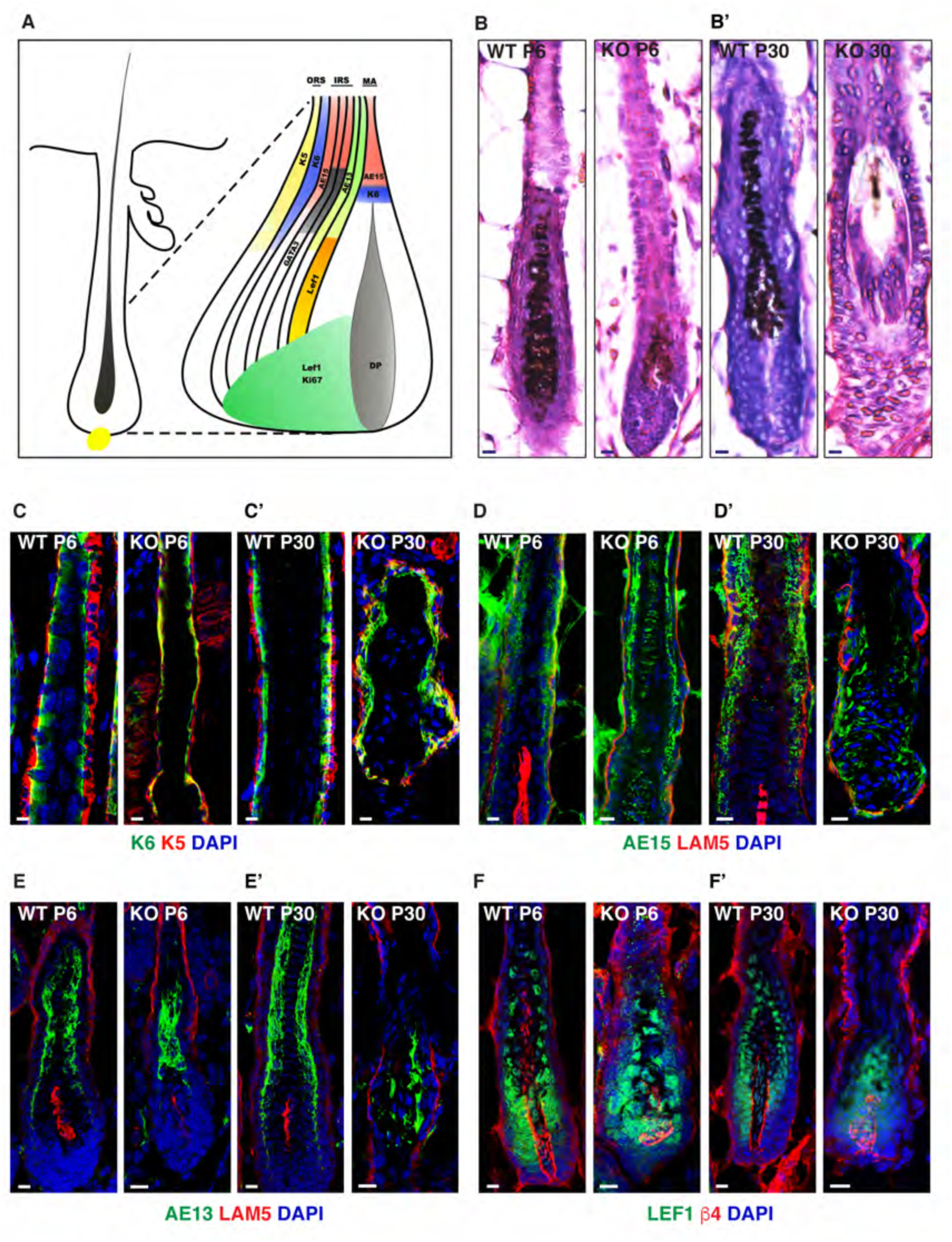
Analysis of the differentiation status of hair follicles in morphogenesis and anagen. Schematic representation of the markers expressed in differentiated layers of hair follicle (A). Histology of P6 and P30 WT and KO follicles (B, B’). Expression of K6 (green), K5 (red) and DAPI (blue) in WT and vinculin KO follicles at P6 and P30 (C, C’). Expression of AE15 (green), laminin 5 (red) and DAPI (blue) in WT and vinculin KO follicles at P6 and P30 (D, D’). Expression of AE13 (green), laminin 5 (red) and DAPI (blue) in WT and vinculin KO follicles at P6 and P30 (E, E’). Expression of Lef1 (green), β4-integrin (red) and DAPI (blue) in WT and vinculin KO follicles at P6 and P30 (F, F’). Scale bar: 10 μM (B-F’).

**Extended supplementary Fig 3:**
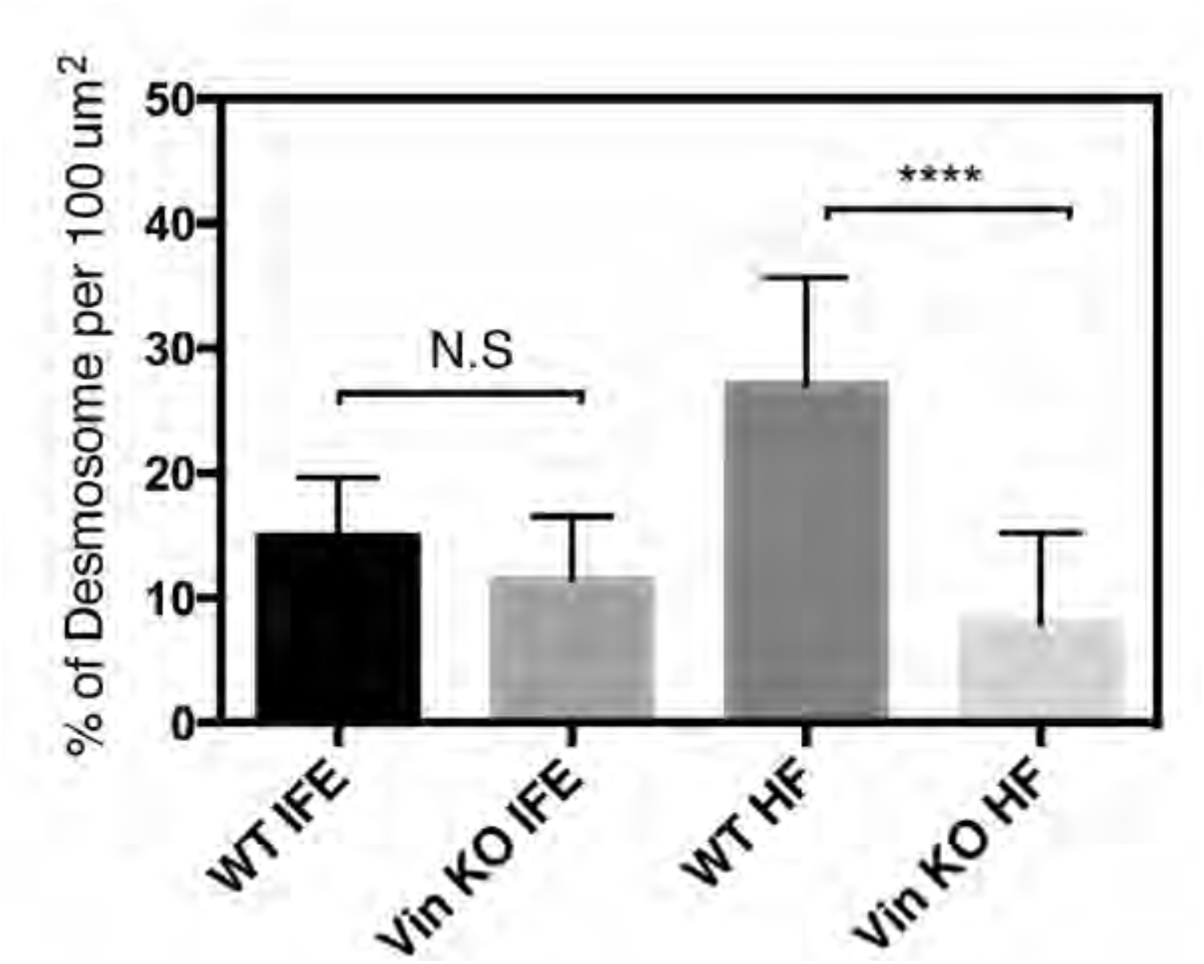
Histogram representing the percentage of desmosomes per 100μm^2^ WT and vinculin KO IFE and hair follicles. \*\*\*\**P* < 0.0001, N.S. not significant.

**Supplementary table 4:**
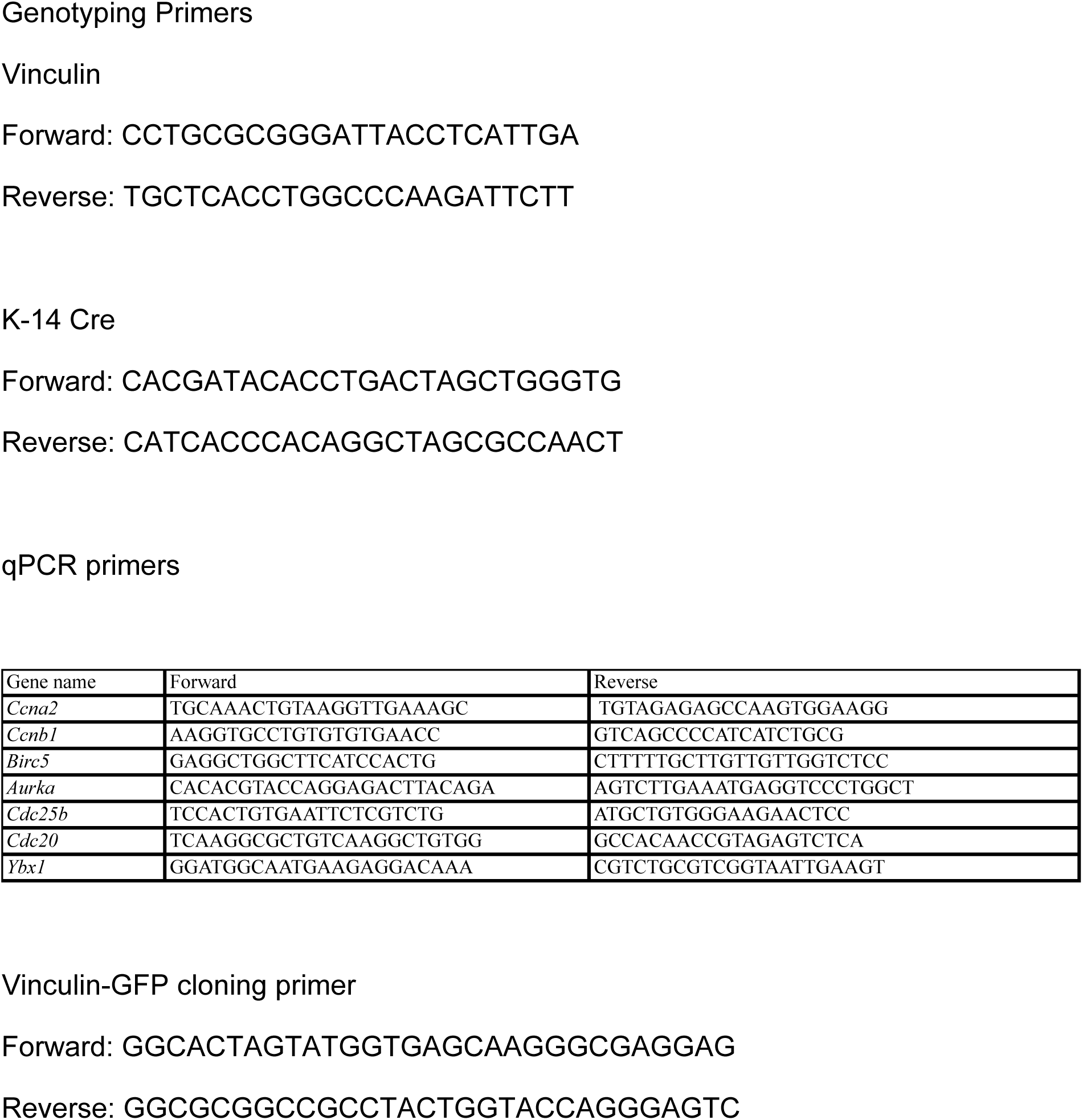
List of primers used for genotyping, qPCR and cloning.

## References

Alatortsev, V.E., Kramerova, I.A., Frolov, M.V., Lavrov, S.A., and Westphal, E.D. (1997). Vinculin gene is non-essential in Drosophila melanogaster. FEBS letters 413, 197–201.

Alenghat, F.J., Fabry, B., Tsai, K.Y., Goldmann, W.H., and Ingber, D.E. (2000). Analysis of cell mechanics in single vinculin-deficient cells using a magnetic tweezer. Biochemical and biophysical research communications 277, 93–99.

Barstead, R.J., and Waterston, R.H. (1989). The basal component of the nematode dense-body is vinculin. Journal of Biological Chemistry 264, 10177–10185.

Bays, J.L., and DeMali, K.A. (2017). Vinculin in cell–cell and cell–matrix adhesions. Cellular and Molecular Life Sciences 74, 2999–3009.

Biggs, L.C., Kim, C.S., Miroshnikova, Y.A., and Wickström, S.A. (2019). Mechanical Forces in the Skin: Roles in Tissue Architecture, Stability, and Function. Journal of Investigative Dermatology.

Blanpain, C., Lowry, W.E., Geoghegan, A., Polak, L., and Fuchs, E. (2004). Self-renewal, multipotency, and the existence of two cell populations within an epithelial stem cell niche. Cell 118, 635–648.

Bolger, A.M., Lohse, M., and Usadel, B. (2014). Trimmomatic: a flexible trimmer for Illumina sequence data. Bioinformatics 30, 2114–2120.

Buckley, C.D., Tan, J., Anderson, K.L., Hanein, D., Volkmann, N., Weis, W.I., Nelson, W.J., and Dunn, A.R. (2014). The minimal cadherin-catenin complex binds to actin filaments under force. Science 346, 1254211.

Cotsarelis, G., Sun, T.-T., and Lavker, R.M. (1990). Label-retaining cells reside in the bulge area of pilosebaceous unit: implications for follicular stem cells, hair cycle, and skin carcinogenesis. Cell 61, 1329–1337.

DasGupta, R., and Fuchs, E. (1999). Multiple roles for activated LEF/TCF transcription complexes during hair follicle development and differentiation. Development 126, 4557–4568.

Fernandez-Sanchez, M.-E., Brunet, T., Röper, J.-C., and Farge, E. (2015). Mechanotransduction’s impact on animal development, evolution, and tumorigenesis. Annual review of cell and developmental biology 31, 373–397.

Ferreira, T., and Rasband, W. (2012). ImageJ user guide. ImageJ/Fiji 1, 155–161.

Gattazzo, F., Urciuolo, A., and Bonaldo, P. (2014). Extracellular matrix: a dynamic microenvironment for stem cell niche. Biochimica et Biophysica Acta (BBA)-General Subjects 1840, 2506–2519.

Gilmour, D., Rembold, M., and Leptin, M. (2017). From morphogen to morphogenesis and back. Nature 541, 311–320.

Goff, L., Trapnell, C., and Kelley, D. (2013). cummeRbund: Analysis, exploration, manipulation, and visualization of Cufflinks high-throughput sequencing data. R package version 2.

Goldmann, W.H. (2016). Role of vinculin in cellular mechanotransduction. Cell biology international 40, 241–256.

Han, M.K., Van Der Krogt, G.N., and De Rooij, J. (2017). Zygotic vinculin is not essential for embryonic development in zebrafish. PloS one 12, e0182278.

Heisenberg, C.-P., and Bellaïche, Y. (2013). Forces in tissue morphogenesis and patterning. Cell 153, 948–962.

Hirai, Y., Nose, A., Kobayashi, S., and Takeichi, M. (1989). Expression and role of E- and P-cadherin adhesion molecules in embryonic histogenesis. II. Skin morphogenesis. Development 105, 271–277.

Hsu, Y.-C., Pasolli, H.A., and Fuchs, E. (2011). Dynamics between stem cells, niche, and progeny in the hair follicle. Cell 144, 92–105.

Huang, D.L., Bax, N.A., Buckley, C.D., Weis, W.I., and Dunn, A.R. (2017). Vinculin forms a directionally asymmetric catch bond with F-actin. Science 357, 703–706.

Huveneers, S., Oldenburg, J., Spanjaard, E., van der Krogt, G., Grigoriev, I., Akhmanova, A., Rehmann, H., and de Rooij, J. (2012). Vinculin associates with endothelial VE-cadherin junctions to control force-dependent remodeling. The Journal of cell biology 196, 641–652.

Imamura, Y., Itoh, M., Maeno, Y., Tsukita, S., and Nagafuchi, A. (1999). Functional domains of α-catenin required for the strong state of cadherin-based cell adhesion. The Journal of cell biology 144, 1311–1322.

Jaks, V., Barker, N., Kasper, M., Van Es, J.H., Snippert, H.J., Clevers, H., and Toftgård, R. (2008). Lgr5 marks cycling, yet long-lived, hair follicle stem cells. Nature genetics 40, 1291.

Jensen, K.B., Collins, C.A., Nascimento, E., Tan, D.W., Frye, M., Itami, S., and Watt, F.M. (2009). Lrig1 expression defines a distinct multipotent stem cell population in mammalian epidermis. Cell stem cell 4, 427–439.

Kim, D., Langmead, B., and Salzberg, S.L. (2015). HISAT: a fast spliced aligner with low memory requirements. Nature methods 12, 357.

Kobielak, K., Pasolli, H.A., Alonso, L., Polak, L., and Fuchs, E. (2003). Defining BMP functions in the hair follicle by conditional ablation of BMP receptor IA. J Cell Biol 163, 609–623.

Lay, K., Kume, T., and Fuchs, E. (2016). FOXC1 maintains the hair follicle stem cell niche and governs stem cell quiescence to preserve long-term tissue-regenerating potential. Proceedings of the National Academy of Sciences 113, E1506–E1515.

Le Duc, Q., Shi, Q., Blonk, I., Sonnenberg, A., Wang, N., Leckband, D., and De Rooij, J. (2010). Vinculin potentiates E-cadherin mechanosensing and is recruited to actin-anchored sites within adherens junctions in a myosin II–dependent manner. The Journal of cell biology 189, 1107–1115.

Le, S., Yu, M., and Yan, J. (2019). Direct single-molecule quantification reveals unexpectedly high mechanical stability of vinculin—talin/α-catenin linkages. Science Advances 5, eaav2720.

Lewis, J.E., Wahl, J.K., Sass, K.M., Jensen, P.J., Johnson, K.R., and Wheelock, M.J. (1997). Cross-talk between adherens junctions and desmosomes depends on plakoglobin. The Journal of cell biology 136, 919–934.

Li, J., Gao, E., Vite, A., Yi, R., Gomez, L., Goossens, S., Van Roy, F., and Radice, G.L. (2015). Alpha-catenins control cardiomyocyte proliferation by regulating Yap activity. Circulation research 116, 70–79.

Liu, Y., Lyle, S., Yang, Z., and Cotsarelis, G. (2003). Keratin 15 promoter targets putative epithelial stem cells in the hair follicle bulge. Journal of Investigative Dermatology 121, 963–968.

Low, B.C., Pan, C.Q., Shivashankar, G., Bershadsky, A., Sudol, M., and Sheetz, M. (2014). YAP/TAZ as mechanosensors and mechanotransducers in regulating organ size and tumor growth. FEBS letters 588, 2663–2670.

Lyon, R.C., Zanella, F., Omens, J.H., and Sheikh, F. (2015). Mechanotransduction in cardiac hypertrophy and failure. Circulation research 116, 1462–1476.

Manabe, M., Mizoguchi, M., Niwa, M., Bertolino, A.P., Ishidoh, K., Kominami, E., and Ogawa, H. (1996). Assembly of hair keratins in transfected epithelial cells. Biochemical and biophysical research communications 229, 965–973.

Martino, F., Perestrelo, A.R., Vinarský, V., Pagliari, S., and Forte, G. (2018). Cellular mechanotransduction: from tension to function. Frontiers in physiology 9.

Ming, P.S. (2019). Mechanosesitive heart functions of aT-Catenin and Cardiac titin.

Miroshnikova, Y.A., Le, H.Q., Schneider, D., Thalheim, T., Rübsam, M., Bremicker, N., Polleux, J., Kamprad, N., Tarantola, M., and Wang, I. (2018). Adhesion forces and cortical tension couple cell proliferation and differentiation to drive epidermal stratification. Nature cell biology 20, 69.

Miyake, Y., Inoue, N., Nishimura, K., Kinoshita, N., Hosoya, H., and Yonemura, S. (2006). Actomyosin tension is required for correct recruitment of adherens junction components and zonula occludens formation. Experimental cell research 312, 1637–1650.

Mootha, V.K., Lindgren, C.M., Eriksson, K.-F., Subramanian, A., Sihag, S., Lehar, J., Puigserver, P., Carlsson, E., Ridderstråle, M., and Laurila, E. (2003). PGC-1α-responsive genes involved in oxidative phosphorylation are coordinately downregulated in human diabetes. Nature genetics 34, 267.

Morgner, J., Ghatak, S., Jakobi, T., Dieterich, C., Aumailley, M., and Wickström, S.A. (2015). Integrin-linked kinase regulates the niche of quiescent epidermal stem cells. Nature communications 6, 8198.

Morris, R.J., Bortner, C.D., Cotsarelis, G., Reece, J.M., Trempus, C.S., Faircloth, R.S., and Tennant, R.W. (2003). Enrichment for living murine keratinocytes from the hair follicle bulge with the cell surface marker CD34. Journal of Investigative Dermatology 120, 501–511.

Morris, R.J., Liu, Y., Marles, L., Yang, Z., Trempus, C., Li, S., Lin, J.S., Sawicki, J.A., and Cotsarelis, G. (2004). Capturing and profiling adult hair follicle stem cells. Nature biotechnology 22, 411.

Müller-Röver, S., Foitzik, K., Paus, R., Handjiski, B., van der Veen, C., Eichmüller, S., McKay, I.A., and Stenn, K.S. (2001). A comprehensive guide for the accurate classification of murine hair follicles in distinct hair cycle stages. Journal of Investigative Dermatology 117, 3–15.

Müller-Röver, S., Tokura, Y., Welker, P., Furukawa, F., Wakita, H., Takigawa, M., and Paus, R. (1999). E-and P-cadherin expression during murine hair follicle morphogenesis and cycling. Experimental dermatology 8, 237–246.

Nagafuchi, A., and Tsukita, S. (1994). The Loss of the Expression of α Catenin, the 102 kD Cadherin Associated Protein, in Central Nervous Tissues during Development: (α catenin/cadherin/cell adhesion/CNS). Development, growth & differentiation 36, 59–71.

Nanba, D., Hieda, Y., and Nakanishi, Y. (2000). Remodeling of desmosomal and hemidesmosomal adhesion systems during early morphogenesis of mouse pelage hair follicles. Journal of investigative dermatology 114, 171–177.

Niessen, C.M., and Gottardi, C.J. (2008). Molecular components of the adherens junction. Biochimica et Biophysica Acta (BBA)-Biomembranes 1778, 562–571.

Nowak, J.A., Polak, L., Pasolli, H.A., and Fuchs, E. (2008). Hair follicle stem cells are specified and function in early skin morphogenesis. Cell stem cell 3, 33–43.

Ohmori, T., Kashiwakura, Y., Ishiwata, A., Madoiwa, S., Mimuro, J., Furukawa, Y., and Sakata, Y. (2010). Vinculin is indispensable for repopulation by hematopoietic stem cells, independent of integrin function. Journal of Biological Chemistry 285, 31763–31773.

Oliver, R.F., and Jahoda, C.A. (1988). Dermal-epidermal interactions. Clinics in dermatology 6, 74–82.

Page, M.E., Lombard, P., Ng, F., Göttgens, B., and Jensen, K.B. (2013). The epidermis comprises autonomous compartments maintained by distinct stem cell populations. Cell stem cell 13, 471–482.

Pang, S.M., Le, S., Kwiatkowski, A.V., and Yan, J. (2019). Mechanical stability of αT-catenin and its activation by force for vinculin binding. Molecular biology of the cell 30, 1930–1937.

Pennings, S., Liu, K.J., and Qian, H. (2018). The Stem Cell Niche: Interactions between Stem Cells and Their Environment. Stem cells international 2018.

Perret, E., Leung, A., Feracci, H., and Evans, E. (2004). Trans-bonded pairs of E-cadherin exhibit a remarkable hierarchy of mechanical strengths. Proceedings of the National Academy of Sciences 101, 16472–16477.

Petridou, N.I., Spiro, Z., and Heisenberg, C.-P. (2017). Multiscale force sensing in development. Nature cell biology 19, 581–588.

Pinheiro, D., and Bellaïche, Y. (2018). Mechanical force-driven adherens junction remodeling and epithelial dynamics. Developmental cell 47, 3–19.

Raghavan, S., Vaezi, A., and Fuchs, E. (2003). A role for αβ1 integrins in focal adhesion function and polarized cytoskeletal dynamics. Developmental cell 5, 415–427.

Rakshit, S., Zhang, Y., Manibog, K., Shafraz, O., and Sivasankar, S. (2012). Ideal, catch, and slip bonds in cadherin adhesion. Proceedings of the National Academy of Sciences 109, 18815–18820.

Rishikaysh, P., Dev, K., Diaz, D., Qureshi, W.M.S., Filip, S., and Mokry, J. (2014). Signaling involved in hair follicle morphogenesis and development. International journal of molecular sciences 15, 1647–1670.

Rognoni, E., and Walko, G. (2019). The Roles of YAP/TAZ and the Hippo Pathway in Healthy and Diseased Skin. Cells 8, 411.

Rübsam, M., Broussard, J.A., Wickström, S.A., Nekrasova, O., Green, K.J., and Niessen, C.M. (2018). Adherens junctions and desmosomes coordinate mechanics and signaling to orchestrate tissue morphogenesis and function: an evolutionary perspective. Cold Spring Harbor perspectives in biology 10, a029207.

Sarpal, R., Yan, V., Kazakova, L., Sheppard, L., Yu, J.C., Fernandez-Gonzalez, R., and Tepass, U. (2019). Role of α-Catenin and its mechanosensing properties in regulating Hippo/YAP-dependent tissue growth. PLoS genetics 15.

Schepeler, T., Page, M.E., and Jensen, K.B. (2014). Heterogeneity and plasticity of epidermal stem cells. Development 141, 2559–2567.

Schindelin, J., Arganda-Carreras, I., Frise, E., Kaynig, V., Longair, M., Pietzsch, T., Preibisch, S., Rueden, C., Saalfeld, S., and Schmid, B. (2012). Fiji: an open-source platform for biological-image analysis. Nature methods 9, 676.

Schlegelmilch, K., Mohseni, M., Kirak, O., Pruszak, J., Rodriguez, J.R., Zhou, D., Kreger, B.T., Vasioukhin, V., Avruch, J., and Brummelkamp, T.R. (2011). Yap1 acts downstream of α-catenin to control epidermal proliferation. Cell 144, 782–795.

Seddiki, R., Narayana, G.H.N.S., Strale, P.-O., Balcioglu, H.E., Peyret, G., Yao, M., Le, A.P., Teck Lim, C., Yan, J., and Ladoux, B. (2018). Force-dependent binding of vinculin to α-catenin regulates cell–cell contact stability and collective cell behavior. Molecular biology of the cell 29, 380–388.

Sheikh, F., Ross, R.S., and Chen, J. (2009). Cell-cell connection to cardiac disease. Trends in cardiovascular medicine 19, 182–190.

Silvis, M.R., Kreger, B.T., Lien, W.-H., Klezovitch, O., Rudakova, G.M., Camargo, F.D., Lantz, D.M., Seykora, J.T., and Vasioukhin, V. (2011). α-catenin is a tumor suppressor that controls cell accumulation by regulating the localization and activity of the transcriptional coactivator Yap1. Sci Signal 4, ra33–ra33.

Spanjaard, E., and de Rooij, J. (2013). Mechanotransduction: vinculin provides stability when tension rises. Current Biology 23, R159–R161.

Subramanian, A., Tamayo, P., Mootha, V.K., Mukherjee, S., Ebert, B.L., Gillette, M.A., Paulovich, A., Pomeroy, S.L., Golub, T.R., and Lander, E.S. (2005). Gene set enrichment analysis: a knowledge-based approach for interpreting genome-wide expression profiles. Proceedings of the National Academy of Sciences 102, 15545–15550.

Sumida, G.M., Tomita, T.M., Shih, W., and Yamada, S. (2011). Myosin II activity dependent and independent vinculin recruitment to the sites of E-cadherin-mediated cell-cell adhesion. BMC cell biology 12, 48.

Swaminathan, V., Alushin, G.M., and Waterman, C.M. (2017). Mechanosensation: a catch bond that only hooks one way. Current Biology 27, R1158–R1160.

Taylor, G., Lehrer, M.S., Jensen, P.J., Sun, T.-T., and Lavker, R.M. (2000). Involvement of follicular stem cells in forming not only the follicle but also the epidermis. Cell 102, 451–461.

Trapnell, C., Hendrickson, D.G., Sauvageau, M., Goff, L., Rinn, J.L., and Pachter, L. (2013). Differential analysis of gene regulation at transcript resolution with RNA-seq. Nature biotechnology 31, 46.

Trapnell, C., Williams, B.A., Pertea, G., Mortazavi, A., Kwan, G., Van Baren, M.J., Salzberg, S.L., Wold, B.J., and Pachter, L. (2010). Transcript assembly and quantification by RNA-Seq reveals unannotated transcripts and isoform switching during cell differentiation. Nature biotechnology 28, 511.

Tumbar, T., Guasch, G., Greco, V., Blanpain, C., Lowry, W.E., Rendl, M., and Fuchs, E. (2004). Defining the epithelial stem cell niche in skin. Science 303, 359–363.

Vasioukhin, V., Bauer, C., Degenstein, L., Wise, B., and Fuchs, E. (2001). Hyperproliferation and defects in epithelial polarity upon conditional ablation of α-catenin in skin. Cell 104, 605–617.

Vasioukhin, V., Bauer, C., Yin, M., and Fuchs, E. (2000). Directed actin polymerization is the driving force for epithelial cell–cell adhesion. Cell 100, 209–219.

Vasquez, C.G., and Martin, A.C. (2016). Force transmission in epithelial tissues. Developmental Dynamics 245, 361–371.

Vidal, V.P., Chaboissier, M.-C., Lützkendorf, S., Cotsarelis, G., Mill, P., Hui, C.-C., Ortonne, N., Ortonne, J.-P., and Schedl, A. (2005). Sox9 is essential for outer root sheath differentiation and the formation of the hair stem cell compartment. Current Biology 15, 1340–1351.

Wang, L.D., and Wagers, A.J. (2011). Dynamic niches in the origination and differentiation of haematopoietic stem cells. Nature reviews Molecular cell biology 12, 643–655.

Watabe-Uchida, M., Uchida, N., Imamura, Y., Nagafuchi, A., Fujimoto, K., Uemura, T., Vermeulen, S., Van Roy, F., Adamson, E.D., and Takeichi, M. (1998). α-Catenin-vinculin interaction functions to organize the apical junctional complex in epithelial cells. The Journal of cell biology 142, 847–857.

Weiss, E.E., Kroemker, M., Rüdiger, A.-H., Jockusch, B.M., and Rüdiger, M. (1998). Vinculin is part of the cadherin–catenin junctional complex: complex formation between α-catenin and vinculin. The Journal of cell biology 141, 755–764.

Wickham, H. (2016). Ggplot2: Elegant graphics for data analysis.,(Springer-Verlag: New York) (USA).

Xu, W., Baribault, H., and Adamson, E.D. (1998). Vinculin knockout results in heart and brain defects during embryonic development. Development 125, 327–337.

Yao, M., Qiu, W., Liu, R., Efremov, A.K., Cong, P., Seddiki, R., Payre, M., Lim, C.T., Ladoux, B., and Mege, R.-M. (2014). Force-dependent conformational switch of α-catenin controls vinculin binding. Nature communications 5, 4525.

Yonemura, S., Wada, Y., Watanabe, T., Nagafuchi, A., and Shibata, M. (2010). α-Catenin as a tension transducer that induces adherens junction development. Nature cell biology 12, 533.

Yu, M., Yuan, X., Lu, C., Le, S., Kawamura, R., Efremov, A.K., Zhao, Z., Kozlov, M.M., Sheetz, M., and Bershadsky, A. (2017). mDia1 senses both force and torque during F-actin filament polymerization. Nature communications 8, 1650.

Zemljic-Harpf, A.E., Miller, J.C., Henderson, S.A., Wright, A.T., Manso, A.M., Elsherif, L., Dalton, N.D., Thor, A.K., Perkins, G.A., and McCulloch, A.D. (2007). Cardiac-myocyte-specific excision of the vinculin gene disrupts cellular junctions, causing sudden death or dilated cardiomyopathy. Molecular and cellular biology 27, 7522–7537.

Zhao, B., Wei, X., Li, W., Udan, R.S., Yang, Q., Kim, J., Xie, J., Ikenoue, T., Yu, J., and Li, L. (2007). Inactivation of YAP oncoprotein by the Hippo pathway is involved in cell contact inhibition and tissue growth control. Genes & development 21, 2747–2761.

